# Systematic Bayesian Posterior Analysis Guided by Kullback-Leibler Divergence Facilitates Hypothesis Formation

**DOI:** 10.1101/2022.07.14.500054

**Authors:** Holly A. Huber, Senta K. Georgia, Stacey D. Finley

## Abstract

Bayesian inference produces a posterior distribution for the parameters and predictions from a mathematical model that can be used to guide the formation of hypotheses; specifically, the posterior may be searched for evidence of alternative model hypotheses, which serves as a starting point for hypothesis formation and model refinement. Previous approaches to search for this evidence are largely qualitative and unsystematic; further, demonstrations of these approaches typically stop at hypothesis formation, leaving the questions they raise unanswered. Here, we introduce a Kullback-Leibler (KL) divergence-based ranking to expedite Bayesian hypothesis formation and investigate the hypotheses it generates, ultimately generating novel, biologically significant insights. Our approach uses KL divergence to rank parameters by how much information they gain from experimental data. Subsequently, rather than searching all model parameters at random, we use this ranking to prioritize examining the posteriors of the parameters that gained the most information from the data for evidence of alternative model hypotheses. We test our approach with two examples, which showcase the ability of our approach to systematically uncover different types of alternative hypothesis evidence. First, we test our KL divergence ranking on an established example of Bayesian hypothesis formation.

Our top-ranked parameter matches the one previously identified to produce alternative hypotheses. In the second example, we apply our ranking in a novel study of a computational model of prolactin-induced JAK2-STAT5 signaling, a pathway that mediates beta cell proliferation. Here, we cluster our KL divergence rankings to select only a subset of parameters to examine for qualitative evidence of alternative hypotheses, thereby expediting hypothesis formation. Within this subset, we find a bimodal posterior revealing two possible ranges for the prolactin receptor degradation rate. We go on to refine the model, incorporating new data and determining which degradation rate is most plausible. Overall, we demonstrate that our approach offers a novel quantitative framework for Bayesian hypothesis formation and use it to produce a novel, biologically-significant insight.

## 1. Introduction

The formation and testing of hypotheses is perhaps the primary endeavor of science. Hypotheses emerge by considering uncertainty in scientific study— if we are completely certain in our knowledge, there is no need to form alternative explanatory hypotheses. Thus, uncertainty is necessary to motivate hypothesis formation. Well-designed experimental studies allow one to formally test hypotheses. For complex biological phenomena, such as intracellular protein signaling networks, mathematical models can produce more quantitative, systems-level insight than experiments alone. Such mathematical models must have accurate parameter values in order to generate reliable predictions. However, common methods of estimating model parameter values may not fully capture uncertainty, and thus overlook alternative hypotheses (Hug et al., 2013; Tötsch and Hoffmann, 2020). Bayesian statistics address this limitation by enabling the mathematical characterization of uncertainty in model parameters and subsequent predictions.

Applied to parameter estimation, Bayesian statistics define the distribution of viable parameters given the available experimental data. Formally, this is known as the *posterior distribution*, the distribution of model parameters conditioned on the data. As informative data are added, the set of viable model parameters changes, and the posterior is updated accordingly. For example, new data might increase our certainty about model parameters and thus cause a decrease in the variance of the posterior distribution. On the other hand, conflicting data might decrease certainty about model parameters, and thus increase the variance of the posterior distribution. If approximated correctly, a posterior can reveal uncertainty in the estimated parameter values, and subsequently, uncertainty in the model predictions. Once this model uncertainty is established, the posterior may be used to formulate new hypotheses.

Indeed, previous studies have demonstrated the utility of posteriors for hypothesis formation based on mechanistic models of protein network dynamics. For example, Bayesian data integration has been used to reveal inconsistencies in a classic study on protease self-digest kinetics (Tötsch and Hoffmann, 2020). Bayesian data integration is the process of integrating subsets of data into the posterior inference and observing whether different data constrain the model in contradictory ways (Thijssen et al., 2018). Using this process, the authors were able to uncover such contradictory information, which inspired an improved explanatory hypothesis for the observed protease dynamics. In another example, qualitative analyses of the Bayesian posterior revealed alternative mechanisms explaining red blood cell signaling (Hug et al., 2013). To do this, the authors first searched each parameter’s posterior for bimodalities.This search revealed two alternative reaction rates that both explained the observed red blood cell signaling response. These alternative rates were then used to characterize testable hypotheses of protein signaling dynamics.

Though both studies successfully demonstrated the utility of posteriors in hypothesis formation, their methods and scope had some limitations. Both approaches—Bayesian data integration and searching for bimodality in the posterior—rely on largely qualitative posterior analyses of all the model parameters, which limits their scalability (Hug et al., 2013; Thijssen et al., 2018). Furthermore, both papers’ scopes were limited to hypothesis formation: the proposed hypotheses remained uninvestigated (Hug et al., 2013; Tötsch and Hoffmann, 2020). Here, we present a systematic posterior analysis approach that facilitates hypothesis formation and, subsequently, investigate the alternative hypotheses it reveals. Our approach uses Kullback-Leibler (KL) divergence to rank parameters by how much information they gain from the experimental data used to infer the posterior distribution. Subsequently, rather than searching all model parameters at random, we use this ranking to prioritize searching the parameters that gained the most information from the data for evidence of alternative model hypotheses. Such evidence may manifest in the posterior, for example, as a bimodal posterior, or as a posterior that is inconsistent with different data subsets, as previously described (Hug et al., 2013; Tötsch and Hoffmann, 2020). Previous work has leveraged KL divergence as a metric of information gained from data to optimize experimental design (Chaloner and Verdinelli, 1995; Lomeli et al., 2021). Here, by using KL divergence to systematically identify parameters whose posterior distribution may indicate alternative hypotheses, we expedite hypothesis formation without needing to know *a priori* how this evidence may manifest in the posterior.

We first test our approach on an established example of Bayesian hypothesis formation, which considers protease self-digest kinetics (Tötsch and Hoffmann, 2020). This example serves to verify our method, while also showing that it can be applied to identify alternative hypotheses that are not reflected as bimodality of the posterior distribution. The majority of the paper presents results from applying our approach in a second example, a novel study of a biologically relevant model of pancreatic beta cell signaling. We choose to study this pathway for its therapeutic potential. Because a decrease in beta cell mass is a hallmark of Type 2 diabetes, previous research has set out to understand signaling pathways that govern beta cell proliferation to design novel approaches to expand functional beta cell mass (Aguayo-Mazzucato and Bonner-Weir, 2018; Brelje et al., 2004; Fujinaka et al., 2007; Jiao et al., 2013; Millette et al., 2022; Rieck et al., 2009; Rønn et al., 2002; Salazar-Petres and Sferruzzi-Perri, 2022). Pregnancy has been a key model system to understand beta cell proliferation.

Observations from both human and rodent pregnancy show that beta cell mass and insulin secretion increase to compensate for increased insulin resistance. These studies revealed prolactin (PRL)-induced janus kinase 2 (JAK2) and signal transducer and activator of transcription 5 (STAT5) signaling as a key regulator of beta cell proliferation (Brelje et al., 2004, 2002). Specifically, the JAK2-STAT5 pathway regulates proliferation by controlling the expression of several proteins. Some of these proteins, such as anti-apoptotic protein B-cell lymphoma-extra large (BCL-xL), influence cell phenotype, while others, such as inhibitor protein Suppressor of Cytokine Signaling (SOCS), form feedback loops with proteins upstream in the pathway to control cell signaling (Fujinaka et al., 2007; Rieck et al., 2009)..

For the first example, we apply our KL divergence ranking directly to the posteriors estimated in Totsch et al. and show how our approach may facilitate their investigation. For the second example, we begin by improving the posterior inference of our previously established model (Mortlock et al., 2021). Subsequently, we analyze the inferred posterior with our KL divergence-based approach, uncovering two plausible receptor degradation rates that both explain the observed SOCS signaling dynamics controlling beta cell proliferation. We then investigate the proposed hypotheses by refining the posterior estimate with new data and validating with literature evidence. Finally, we apply the improved model to resolve the proposed hypotheses, thereby answering an open question in pancreatic beta cell biology: *what are the SOCS negative feedback dynamics controlling PRL-induced JAK2-STAT5 signaling that drive beta cell proliferation?*

## 2. Materials and Methods

### 2.1 KL Divergence Ranking for Prioritizing Parameters that Gain Information from New Data

We use Kullback-Leibler (KL) divergence to rank model parameters by how much information they gain from incorporating experimental data into our posterior inference procedure. KL divergence quantifies the difference between two distributions, where a larger KL divergence corresponds to a larger difference. We use univariate kernel density estimates and Monte Carlo integration to approximate the KL divergence. Further details on this approximation are included in the **Supplementary Materials**.

We calculate the KL divergence from the marginal prior distribution to the marginal posterior distribution. Since the prior is a distribution over the parameters while the posterior is a distribution over the parameters given the experimental data, if there is a difference between the prior and posterior distribution, it can only be because the experimental data influences the posterior distribution. The more the data informs the posterior, the larger the difference between the prior and posterior, and the larger the KL divergence from the prior to the posterior. In this way, the KL divergence from the marginal prior distribution to the marginal posterior distribution provides a quantitative measure for ranking parameters by how much information they gain from the data.

### 2.2 Test Case: Analysis of a Published Study of Protease Self-digest kinetics

In Totsch et al., the authors model protease self-digest kinetics. Initially, they use a model developed in 1981 to describe this system. This ordinary differential equation (ODE) model comprises five kinetic rate constants, whose posterior distribution is estimated via Bayesian inference using two datasets, referred to as D_SAS and D_REG. However, the authors ultimately find that this mathematical model cannot explain both datasets and propose a new model of self-digest kinetics. In our work, we focus on how to expedite Bayesian data integration, the process Totsch et al. uses to discover that the 1981 model is inadequate.

Bayesian data integration refers to the process of sequentially incorporating new data into the posterior inference procedure to understand what information is contained in each data set (Thijssen et al., 2018). By comparing posterior distributions estimated with different data sets, one may see whether data sets from the same experimental setup inform a model posterior in contradictory ways, implying that the current model is inadequate to explain all the available data and that an alternative model hypothesis is needed.

Specifically, the authors separately estimate the posterior distribution for the D_SAS dataset and the posterior for the D_REG dataset. Comparing the marginal posterior distributions between these datasets reveals that one parameter is constrained differently by the different datasets: the kinetic rate constant *k*_*BR*._. To facilitate uncovering this evidence, our KL Divergence ranking should prioritize this parameter over the other parameters for subsequent analysis. Thus, based on the results of Totsch et al, we designate *k*_*BR*_ as the target parameter for our KL Divergence ranking for this test case.

### 2.3 Case Study: Novel Study of PRL-induced Signaling in Pancreatic Beta Cells

#### 2.3.1 Modeling Framework

We modeled PRL-induced intracellular signaling in rodent pancreatic beta cells (**Figure 1**). During pregnancy, PRL induces beta cell proliferation in rodents via the JAK2-STAT5 signaling cascade (Brelje et al., 2004, 2002). Specifically, PRL binds to the prolactin receptor (PRLR), inducing subsequent JAK2 and STAT5 activation (phosphorylation). After activation, STAT5 translocates into the nucleus, where it regulates the transcription of various gene products. STAT5 mediates transcription of proteins that provide positive and negative feedback to the signaling pathway, PRLR and the SOCS inhibitor family, respectively (Rieck et al., 2009). STAT5 also promotes transcription of proteins that characterize a proliferative cell phenotype, such as the anti-apoptotic protein BCL-xL (Fujinaka et al., 2007). Finally, it is important to note that two different isoforms of STAT5, STAT5A and STAT5B, are activated by this cascade. These proteins can form homo- and hetero-dimers, which follow different translocation patterns upon activation (Brelje et al., 2004, 2002).

**Figure 1.**
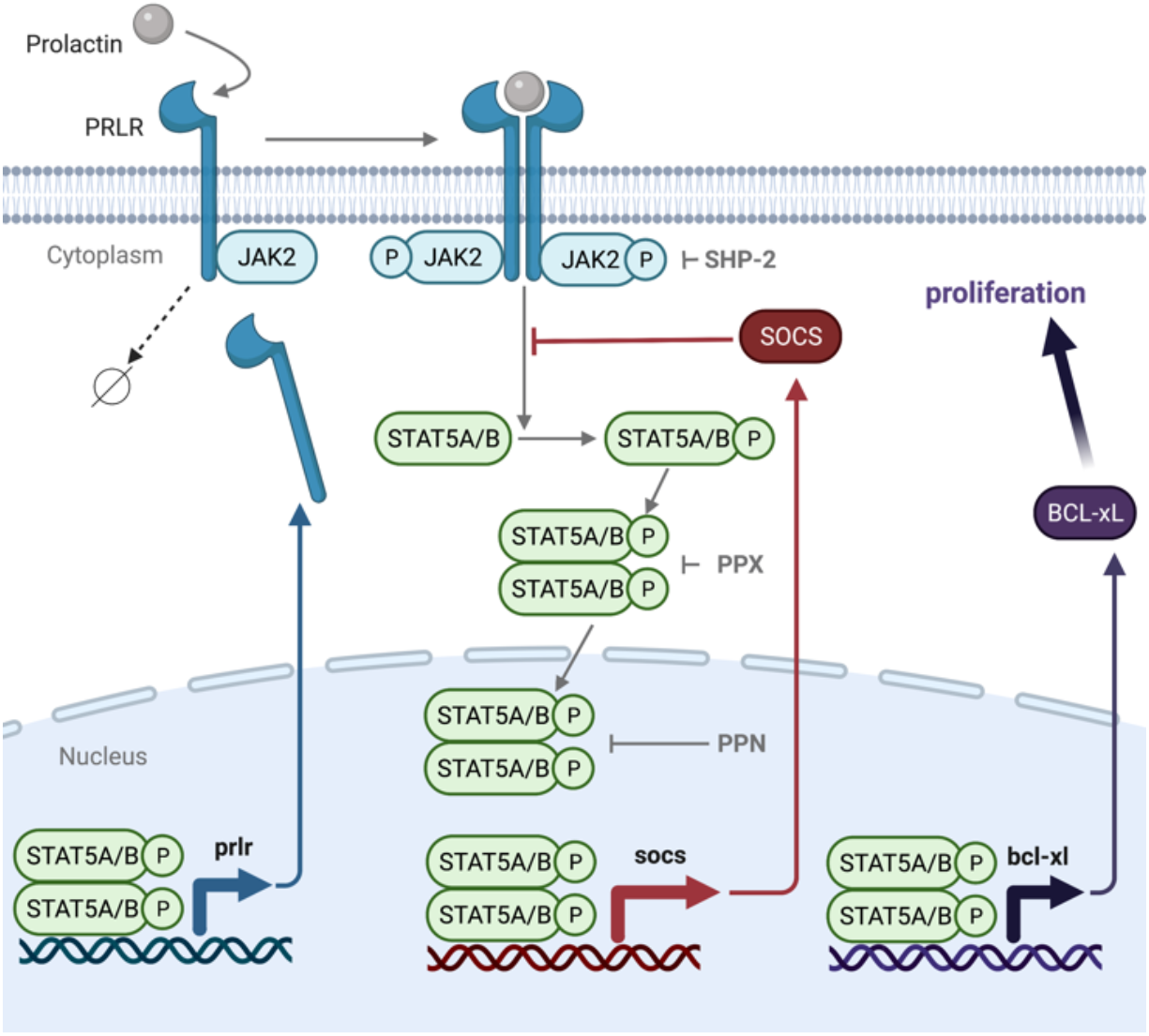
Schematic of the signaling cascade modeled in this work. PRL binds to the PRLR-JAK2 complex, leading to receptor dimerization and JAK2 activation. The activated complex promotes phosphorylation of STAT5A and STAT5B, which dimerize and can translocate to the nucleus. Nuclear STAT5A/B promotes transcription and translation of PRLR, BcL-xL, and SOCS. Feedback from these proteins and the effect of phosphatases (PPX and PPN) are indicated.

To describe this signaling network, we use the ordinary differential equation (ODE) model structure developed by Mortlock et al. This model contains rate equations for each of the described signaling events. In addition, it accounts for degradation of the signaling species, including receptor degradation, as it influences the pathway dynamics. Specifically, the rate of receptor degradation depends on whether PRL is bound or not. Finally, this model also includes rate equations accounting for the effect of nuclear (PPN) and cytoplasmic (PPX) phosphatases observed in the canonical JAK2-STAT5 pathway (Yamada et al., 2003). In total, the model comprises 55 species and 60 kinetic parameters, 33 of which we estimate, as in Mortlock et al. These 33 parameters were selected for model fitting based on either sensitivity-based identifiability analysis results or to include all model submodules (Marino et al., 2008). The rate equations are governed by mass action kinetics or Michaelis-Menten kinetics. This model was most recently used to study the effect of heterogeneity in the intracellular protein concentrations (Simoni et al., 2022). Now, we use it as the basis for this work.

#### 2.3.2 Training Data

Initially, we use the same training data as in Mortlock et al. to estimate our model parameters. Subsequently, we obtained additional data at extended timepoints and refine our parameter estimates using this data. The original training data for our model comprises western blot and immunohistochemical measurements of protein responses in INS-1 cells treated with 200 ng/ml of PRL. Experimental outputs include: (1) phosphorylated JAK2 (pJAK2) up to 6 hours post-PRL stimulation, (2) phosphorylated STAT5A (pSTAT5A) up to 6 hours post-PRL stimulation, (3) phosphorylated STAT5B (pSTAT5B) up to 6 hours post-PRL stimulation, (4) BCL-xL up to 24 hours post-PRL stimulation, (5) the ratio of nuclear to cytoplasmic STAT5A (N:C STAT5A) up to 6 hours post-PRL stimulation, and (6) the ratio of nuclear to cytoplasmic STAT5B (N:C STAT5B) up to 6 hours post-PRL stimulation.

The updated training data for our model includes all the original training data, plus three additional measurements: the relative levels of phosphorylated JAK2, phosphorylated STAT5A, and phosphorylated STAT5B 12 hours post-PRL stimulation. These new timepoints were provided by the authors of the original experimental work (Brelje et al., 2004, 2002). **Figure 2** provides a comparison of the original training data to the updated training data.

**Figure 2.**
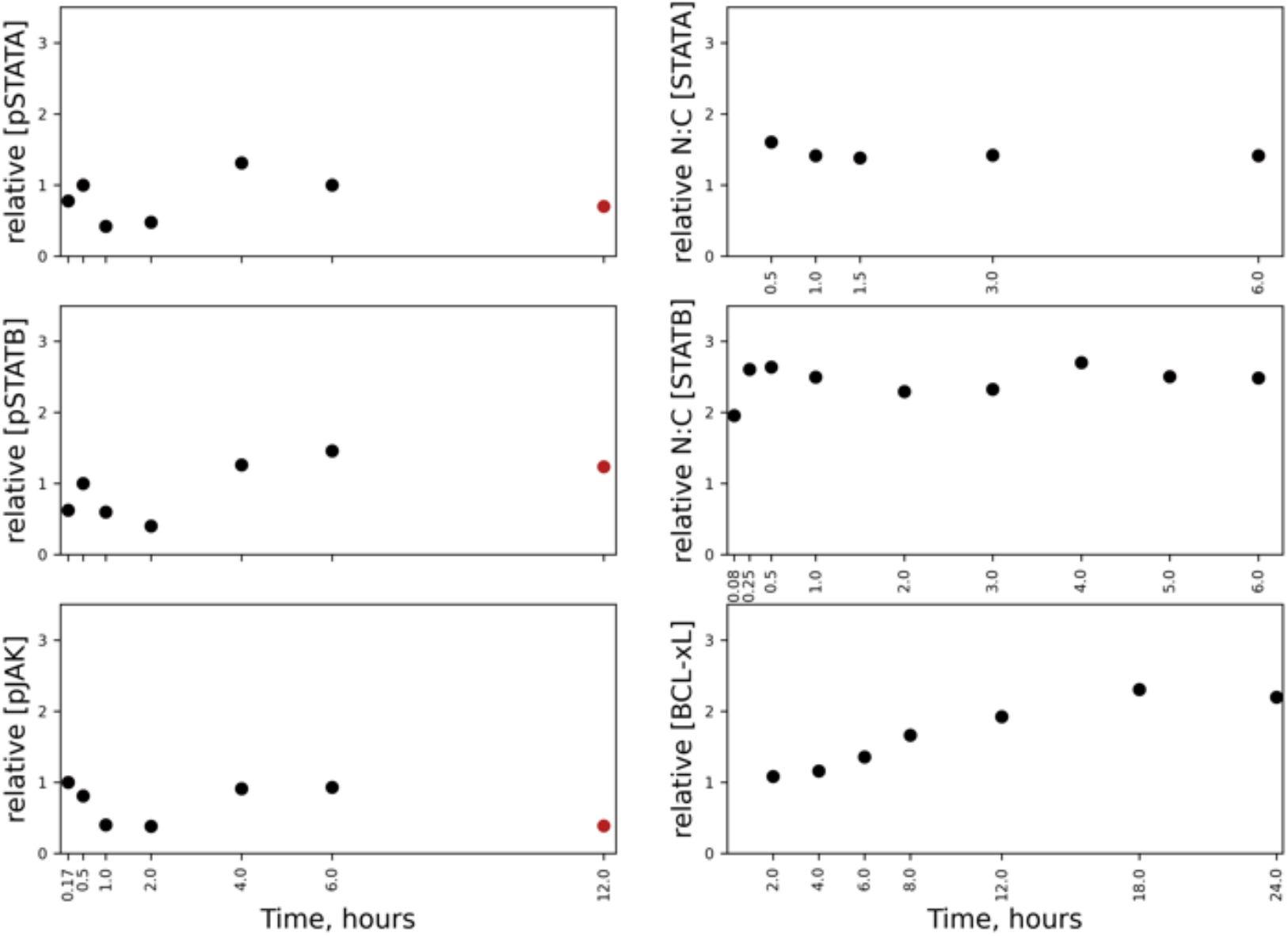
Experimental measurements of proteins included in our model. Measurements are from Brelje (2002) and Fujinaka (2007). Two sets of model training data are displayed—the original set (black), and an extended set, which includes all original data plus three extended time course measurements, which are indicated in red.

#### 2.3.3 Bayesian Posterior Inference

We use the same prior and likelihood distributions as Mortlock et al. to formulate our posterior distribution. **Supplementary Table S1** contains detailed definitions of our prior, likelihood, and proposal distributions. Briefly, the prior distribution is defined as lognormal, centered on the lowest-error parameters found for a model of JAK-STAT signaling in liver cells, with a variance of 2 (Yamada et al., 2003). This variance was assigned based on the variance of kinetic rates pooled across species and pathways (Bar-Even et al., 2011). The likelihood distribution is defined as normal, centered on the data mean with an inverse-gamma distributed variance. The shape of the inverse gamma distribution is defined by two hyperparameters, assigned based on a previously developed posterior estimation procedure (Ghasemi et al., 2011; Makaryan and Finley, 2020). Using these analytical forms for the prior and likelihood distribution, we can approximate the posterior distribution.

We improve upon the posterior approximation procedure from Mortlock et al. by implementing established metrics to monitor the approximation algorithm’s convergence. Like Mortlock et al., we use the Metropolis-Hasting algorithm to estimate the posterior. This algorithm generates an aperiodic and irreducible Markov chain that satisfies the detailed balance condition; samples generated from this Markov chain asymptotically mimic samples drawn from the target distribution, here, the posterior distribution(Andrieu et al., 2003). However, while samples converge to the posterior distribution as the number of draws approaches infinity, a well-known challenge when implementing Markov chain methods is ensuring convergence to the posterior distribution with finite samples (Andrieu et al., 2003; Gelman, 2014).

To address this challenge, we compare samples from at least three independent Markov chains via established qualitative (trace plots, autocorrelation plots, and histograms) and quantitative (split potential scale reduction factor, 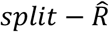) diagnostics (Cowles and Carlin, 1996; Gelman, 2014). Only after samples from these three chains are sufficiently similar 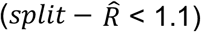 do we conclude convergence and begin analysis of the posterior distribution(Brooks and Gelman, 1998). More details regarding our posterior approximation procedure and the applicability of the 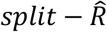 diagnostic to our posterior approximation problem can be found in the **Supplementary Materials**.

#### 2.3.4 K-Means Clustering of KL Divergences to Determine a Subset of Parameters for Investigation

We use Kullback-Leibler (KL) divergence to rank the 33 estimated parameters of the JAK-STAT model by how much information they gain from incorporating experimental data into our posterior inference procedure. Then, we apply K-means clustering to the 33 KL divergences associated with each parameter. We cluster the divergences in order to objectively determine a subset of the 33 parameters for subsequent investigation.

K-means clustering is an unsupervised learning algorithm that partitions data into *k* disjoint sub-groups, or clusters. The objective of this clustering is to group similar data points together and uncover underlying patterns. Further details on this objective are included in the **Supplementary Materials**. We use scikit learn’s *KMeans* function to estimate the optimal group assignment (Pedregosa et al., 2011). We parameterize the *KMeans* function with *k* = 2 clusters. We use 2 clusters because the benefit, that is, increasing similarity, of using more clusters becomes marginal after 2 (**Supplementary Figure S2**). Further details on how we chose 2 clusters is included in the **Supplementary Materials**.

#### 2.3.5 Testing for Multimodality

To compare our approach to a search for multimodality amongst the parameters, we used SciPy’s *find_peaks* function on the kernel density estimates for each marginal posterior distribution (Pedregosa et al., 2011).

#### 2.3.6 Constructing a Validation Metric from Published Experimental Studies

We use the dynamics of SOCS to test plausible biological hypotheses generated via our posterior analysis. We use SOCS as a test output because it provides negative feedback to the JAK-STAT pathway and thus plays a key role in controlling pathway dynamics. Ideally, we would test our model with measurements of the SOCS concentration in beta cells 0-24 hours following PRL stimulation. Although no direct measurements exist, we were able to find indirect, but nevertheless relevant, evidence pertaining to SOCS dynamics in the literature (Chong et al., 2001; Croker et al., 2008; Endo et al., 2003; Jiao et al., 2013; Ram and Waxman, 1999; Rieck et al., 2009). Subsequently, we synthesized this evidence, to create a validation metric.

First, we determined which member(s) of the SOCS family of proteins likely inhibit PRL-induced signaling in beta cells during pregnancy. To do this, we referenced both quantitative PCR (qPCR) data and luciferase assay data. The qPCR data compared the SOCS transcriptional response of rodents between pregnant and non-pregnant states. The data from two such studies confirms that of the eight SOCS proteins, only CISH and SOCS2 are upregulated during rodent pregnancy (**Supplementary Figure S3A and S3B**) (Jiao et al., 2013; Rieck et al., 2009). These results are confirmed by another study that investigated the binding interactions between CISH and PRLR, our pathway’s receptor, via a yeast tri-hybrid system and luciferase assay (Endo et al., 2003). This study found that CISH binds to PRLR and inhibits its activity. Thus, at least CISH has been proven to bind with the receptor in the pathway being modeled here. Based on these three studies, we conclude that either CISH or SOCS2 inhibits PRL-induced JAK2-STAT5 signaling.

Next, we determined the most likely CISH/SOCS2 dynamics from 0-24 hours following ligand stimulation. Although we lacked measurements of this output for PRL-induced JAK2-STAT5 signaling in beta cells, we found consistent CISH/SOCS2 dynamics across other ligands and tissue types: transient SOCS activation, with an early peak (Chong et al., 2001; Ram and Waxman, 1999). Protein dynamics are not necessarily conserved between different ligand and/or tissues (Miller et al., 2019; Nandagopal et al., 2018). However, this consistency across experimental conditions suggests that it is reasonable to assume these same dynamics describe SOCS behavior in our system. To strengthen this assumption, we specifically sought data from beta cells or ligands related to the hormone, PRL.

First, we found an experiment studying CISH, SOCS1, and SOCS2 in beta cells. In this study, beta cells were incubated with either IFN-γ, IFN-α, IL-1β, or TNF-α, and western blot measurements for each of the three SOCS proteins were recorded at 1, 2, 4, and 24 hours post-incubation (Chong et al., 2001). Strikingly, a reliable difference between SOCS1 dynamics and CISH/SOCS2 dynamics emerged across ligands. While SOCS1 exhibited sustained activation 24 hours following stimulation with no early peak, both SOCS2 and CISH exhibited transient activation 24 hours following incubation, with an early peak. **Supplementary Figure S3C** displays a subset of these results.

We next checked that this difference between SOCS1 and CISH/SOCS2 dynamics across different ligands is mechanistically plausible. We found that CISH and SOCS2 share mechanistic similarities, compared to other SOCS proteins like SOCS1. In general, SOCS proteins have three possible inhibition mechanisms (Croker et al., 2008; Ram and Waxman, 1999). Importantly, SOCS2 and CISH share the same mechanism: competitive inhibition with STAT via receptor phosphotyrosine binding, while SOCS1 does not (Croker et al., 2008; Ram and Waxman, 1999). Thus, it is plausible that CISH/SOCS2 would have similar dynamics due to this mechanistic similarity.

Finally, we sought an experiment confirming these consistent dynamics for a ligand more closely related to PRL. We examined published results for CISH dynamics following stimulation with Growth Hormone (GH). Because both GH and PRL are hormones, and GH activates JAK2-STAT signaling, GH-induced SOCS dynamics provide a biologically comparable system to PRL-induced JAK2-STAT5 signaling. In the study, rats were treated with GH and sacrificed 0.5, 1, 1.5, 4, and 24 hours post stimulation, and the livers were harvested for analysis. CISH mRNA expression was measured at each of these timepoints. Again, CISH displayed transient activation by 24 hours following ligand stimulation with an early peak (Ram and Waxman, 1999).

Thus, based on the consistent CISH/SOCS2 dynamics observed across tissues and ligands related to our system of interest, we designate the following as our target output for model validation: *transient SOCS activation 24 hours following stimulation with PRL, with an early peak*.

#### 2.3.7 SOCS Shape Classification for Validation Comparison

To compare our model predictions with the target output, we introduced two shape classification criteria that categorized the predicted SOCS dynamics as: (1) displaying an early peak or no early peak and (2) transient or sustained. We apply these two criteria to the distribution of SOCS predictions generated by simulating the model with the parameter sets sampled from the posterior. In total, there are four possible SOCS shapes: no peak and transient activation, early peak and transient activation, no peak and sustained activation, and early peak and sustained activation. **Supplementary Figure S4** displays examples of each of these four shapes, as well as a graphical representation of our classification criterion. We note that while we apply a discrete shape classification, the transition in the SOCS’ response from one shape to another is somewhere between continuous and discrete: it moves gradually from no peak and transient activation to a large early peak and sustained activation, but with particular mass on either end of this spectrum. To ensure that our discrete classification is suitable to robustly characterize the SOCS’ responses, we varied the classification criterion and checked that our results were conserved, as detailed below.

To categorize SOCS time courses as early peak or no early peak, we applied SciPy’s *find_peaks* function to the first five hours of SOCS dynamics (Pedregosa et al., 2011). The ‘early’ window designation of 0-4 hours was selected based on the CISH/SOCS2 western blot measurements (Chong et al., 2001) (**Supplementary Figure S3C)**. To ensure our results were robust to changes in this window, we also applied the *find_peaks* function to predictions from 0-3 hours. Changing the window did not change the results (**Supplementary Figure S5)**.

Next, to categorize SOCS time courses as transient or sustained, we applied a threshold to the relative SOCS concentration 24 hours following ligand stimulation. A prediction was transient if the SOCS concentration was less than 50% of the maximum SOCS concentration 24 hours post ligand stimulation. We chose the 24 hour time point to be consistent with the western blot data sets, which collected measurements 24 hours post-ligand stimulation (Chong et al., 2001; Ram and Waxman, 1999). The 50% cutoff was derived based on the relative CISH/SOCS2 concentrations collected in the same data sets. To ensure our results were robust to changes in this cutoff, we varied the threshold from 30% to 70%. Changing the threshold did not change the conclusions drawn from the results (**Supplementary Figure S5)**.

#### 2.3.8 Computational Implementation

All code for model analysis is available in the GitHub repository: https://github.com/FinleyLabUSC/KL-Divergence-Bayesian-Hypothesis-Formation.

## 3. Results

### 3.1 Test Case: Analysis of a Published Study of Protease Self-digest kinetics

#### 3.1.1 KL Divergence ranks first the posterior containing alternative hypothesis evidence

Our KL divergence successfully highlights the parameter of interest in our test case, k_BR_. We calculate the divergences between each marginal prior and posterior distributions of the five model parameters calibrated with the D_SAS dataset, and then use these divergences to rank the parameters by how much information they gain from the data (**Figure 3A**). Of the five, parameter k_BR_ has the largest divergence measure (**Figure 3B**) and would therefore be the first parameter selected for further investigation by our method. This selection is in line with the results of Totsch et al., who found that the experimental data sets for their system, D_SAS and D_REG, informed the k_BR_ posterior in contradictory ways. Ultimately, this evidence motivated the authors to develop an alternative model hypothesis. Thus, our KL divergence ranking successfully highlighted the marginal posterior manifesting alternative hypothesis evidence in this established study. This test case provides confidence that our KL divergence-based method can uncover parameters that whose posterior distributions contain evidence of alternative hypotheses.

**Figure 3.**
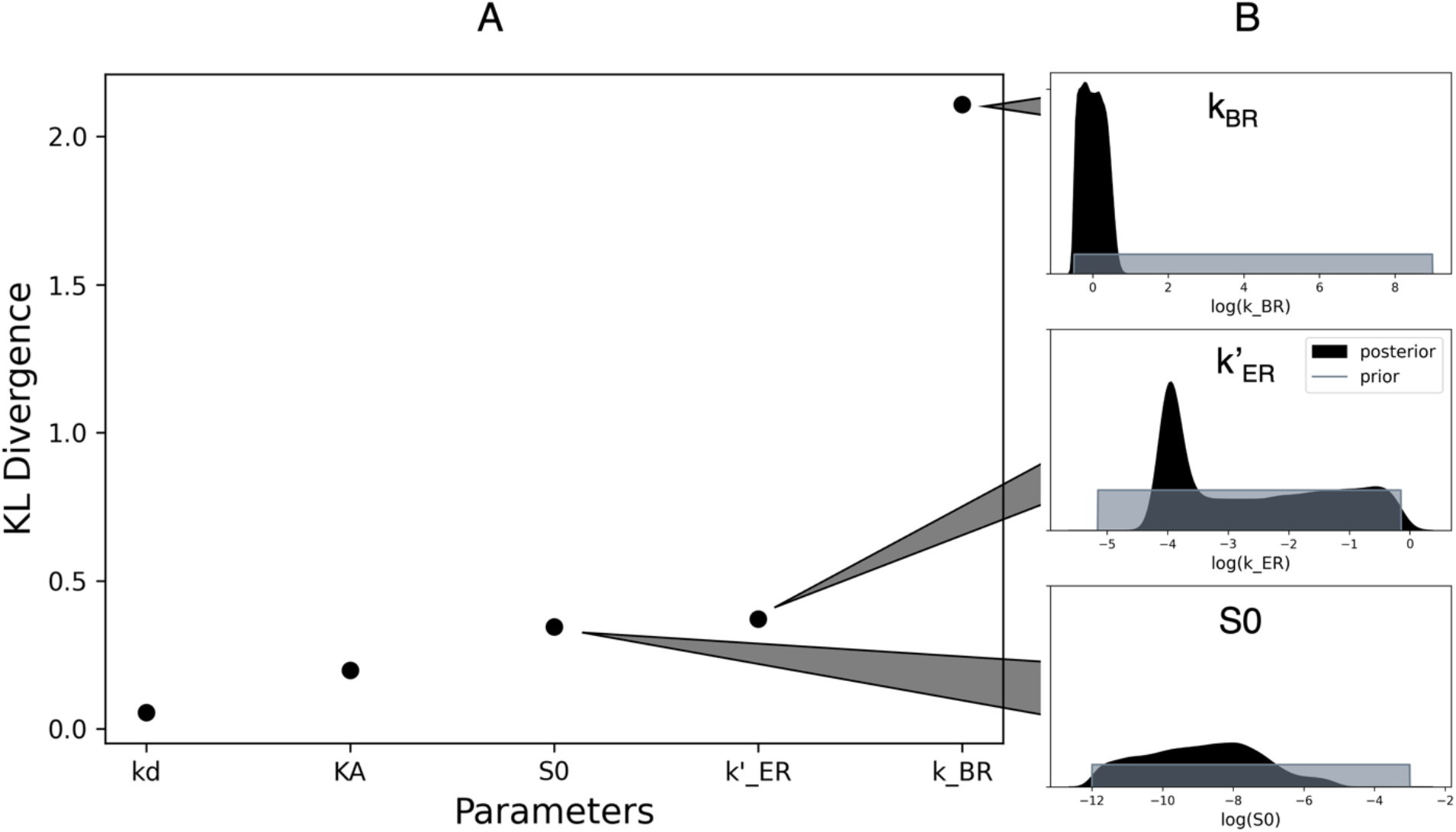
Posterior Analyses for Test Case. (A) Estimated KL divergences from each parameter’s marginal prior distribution to its marginal posterior distribution. (B) Marginal prior and posterior distributions for parameters with the three largest divergences, displayed on a log_10_ scale. Grey, prior; black, posterior.

### 3.2 Case Study: Novel Study of PRL-induced Signaling in Pancreatic Beta Cells

#### 3.2.1 Calibrated Mathematical Model Recapitulates Experimental Data

We established a model of pancreatic beta cell signaling dynamics that is consistent with the experimental data. To begin, the model parameters were fitted via Bayesian inference. We then confirmed that the Bayesian inference method converged to the posterior distribution based on our diagnostic measures (**Supplementary Figure S6A and S7A**). Next, by sampling parameter sets from the posterior distribution, we generated 10,000 predictions of the signaling species’ mean dynamics. These model predictions were subsequently verified against the training data set (**Figure 4**). The majority of training data fall within the 5^th^ to 95^th^ percentile bounds of the model predictions.

**Figure 4.**
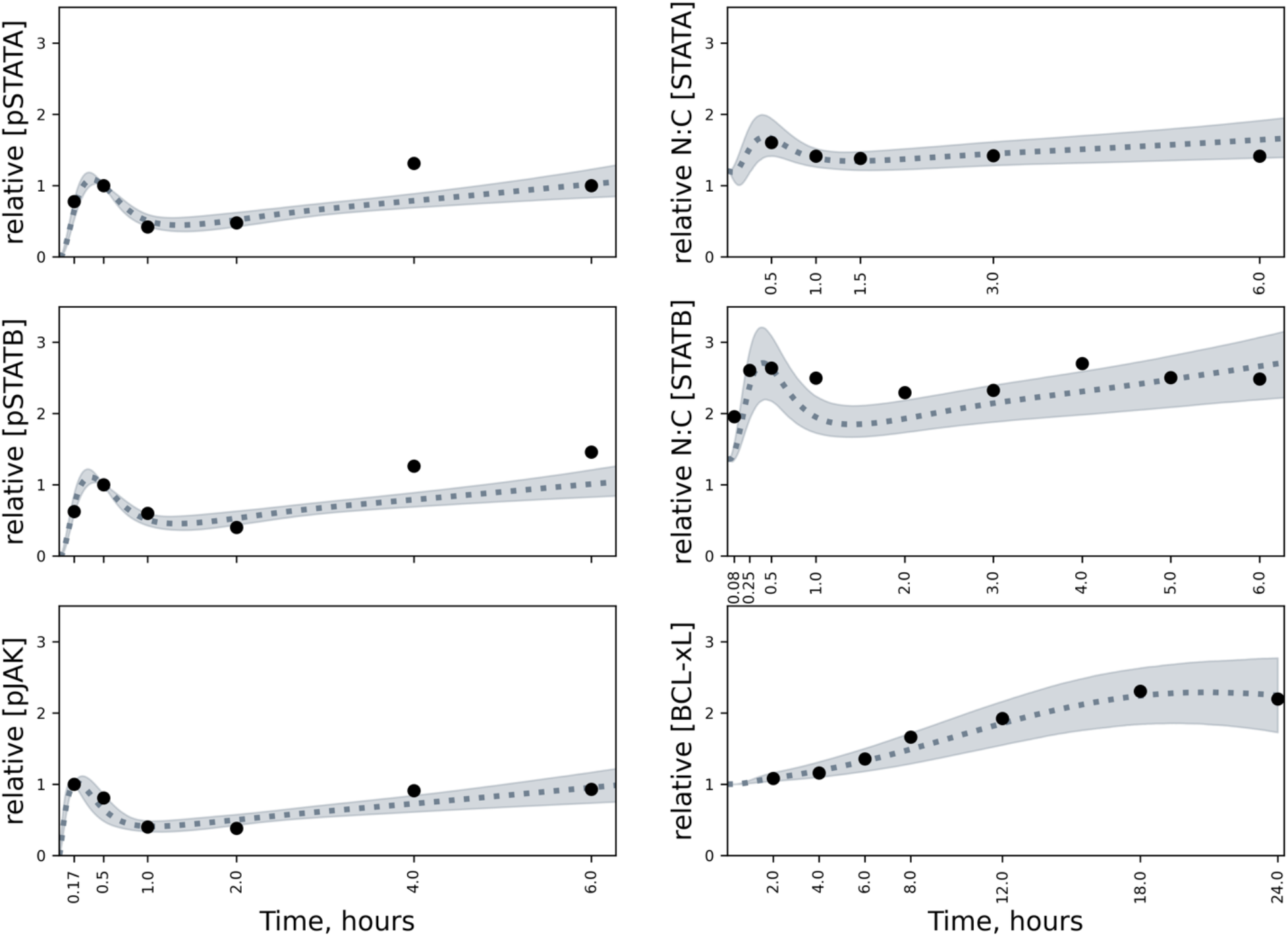
Comparison of inferred model predictions to training data. Predictions of the mean were generated by simulating the model with parameter samples from the posterior distribution. Dashed line, median prediction; shading, the 5^th^ to 95^th^ percentiles of predictions; black dots, experimental data mean.

One systematic deviation between experimental data and our model predictions does exist: the 4-hour time-point for pJAK2, pSTAT5A, pSTAT5B, and N:C STAT5B. Additions to the model could be considered to better recapitulate this time-point; however, as this model acts as a starting point for our subsequent generation, testing, and refinement of hypotheses, we concluded that it sufficiently explained the data and could be used as the basis for model refinement.

#### 3.2.2 Systematic Posterior Analysis Identifies Alternative Biological Hypotheses

Analyses of our inferred posterior distribution identified two plausible PRL:PRLR:SOCS degradation mechanisms driving the signaling response. To quantitatively identify such mechanisms, we first used the KL divergence from the marginal prior distribution to the marginal posterior distribution for each model parameter to rank parameters by how much information they gain from the experimental data (**Figure 5A**). We then used K-means to cluster these rankings into groups to determine any underlying patterns within the divergences. Our data was best described by two clusters (**Supplementary Figure S2**). Interestingly, the three parameters comprising the cluster most informed by the data are all related to PRLR degradation— the degradation rate of PRLR mRNA (*k27*), the degradation rate of the PRL:PRLR complex (*deg_ratio*), and the degradation rate of the PRL:PRLR:SOCS complex (*k24*). The SOCS protein is an inhibitor that provides negative feedback on the signaling pathway. Therefore, when the receptor is bound by SOCS, as in the PRL:PRLR:SOCS complex, PRLR is inactive.

**Figure 5.**
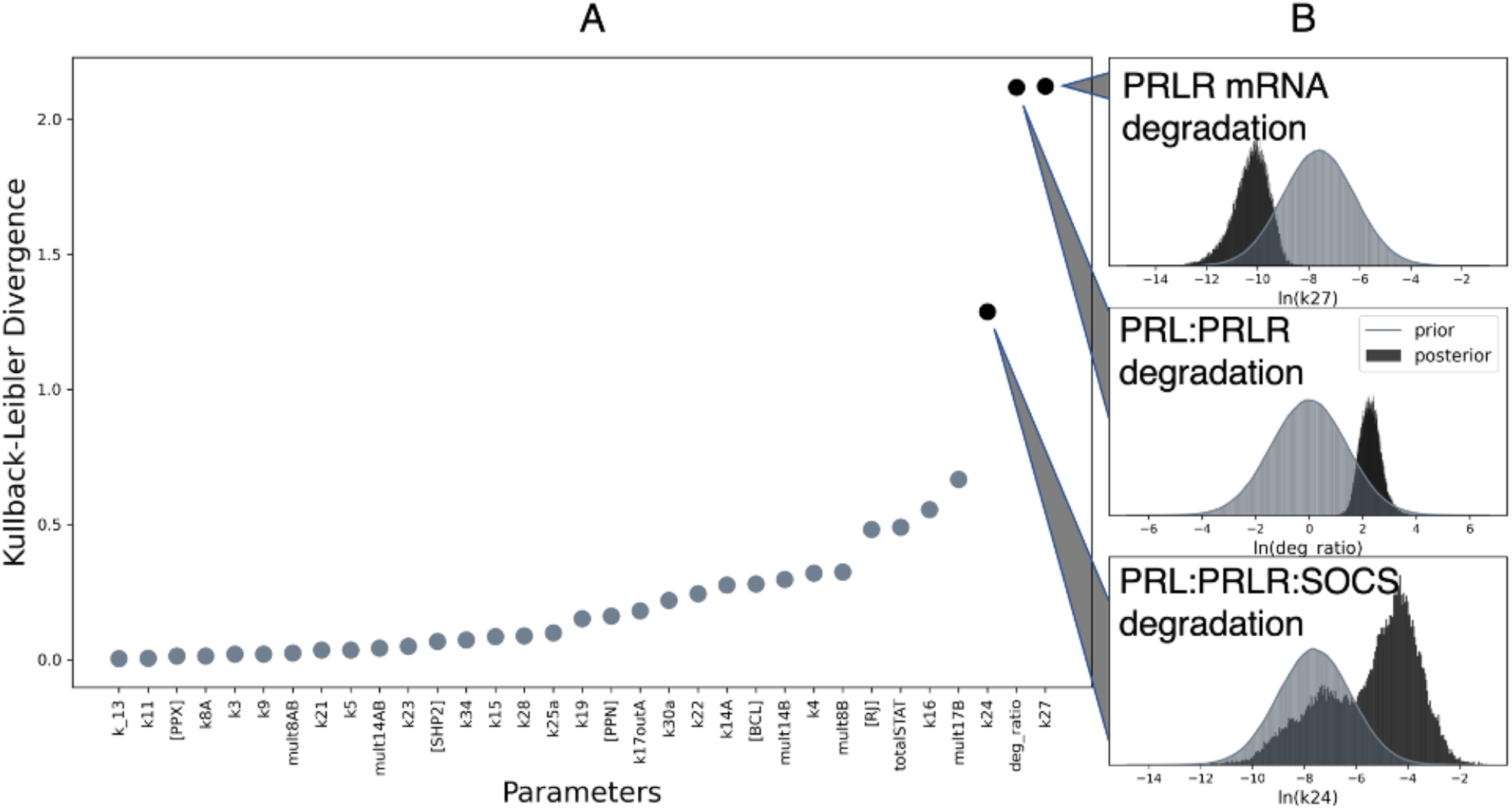
Posterior Analyses for JAK2-STAT5 Case Study. (A) Estimated KL divergences from each parameter’s marginal prior distribution to its marginal posterior distribution with significance cutoff. Grey dots, divergences assigned to cluster 1; black dots, divergences assigned to cluster 2. (B) Marginal prior and posterior distributions for parameters with the largest divergence, displayed on a natural-log scale. Grey, prior; black, posterior.

We next searched for qualitative evidence of divergent mechanisms in the marginal distributions of these top three, data-informed parameters. As one would expect, all three marginal posteriors are observably different from their respective marginal prior distributions (**Figure 5B**). Interestingly, we see that the marginal PRL:PRLR:SOCS complex degradation rate is bimodal, indicating that there are two plausible regimes for the rate at which the SOCS-bound receptor complex is degraded. After uncovering this bimodal distribution via our KL divergence-based clustering, we implemented a multimodality search over the marginal posterior distributions for comparison. This search flagged 12 marginal posterior distributions as multi-modal, many of which contain very little mass under one mode—to the extent that one may consider some of these multi-modal distributions as false positives (**Supplementary Figure S8**). In summary, we use KL divergence to rank parameters by how much information they gain from the data and then search the most data-informed parameters for evidence of alternative model hypotheses in their marginal posterior distribution. This approach uncovered two SOCS-bound receptor degradation regimes that are both plausible given the training data, without *a priori* knowledge of how these alternative regimes would manifest: as a bimodality. Even if we knew that we were searching for multimodal distributions, our approach outperforms a bimodality search, as it returns a smaller subset of parameters for further investigation that also includes the most obviously bimodal distribution amongst all 33 estimated parameters.

Once we uncovered alternative PRL:PRLR:SOCS degradation regimes, we sought to determine if the two possible degradation regimes lead to distinct model predictions. To do this, we first defined two groups of parameter sets from the posterior distribution samples. One group contained 3000 parameter sets with slower PRL:PRLR:SOCS degradation values (**Figure 6A**, blue), while the other group contained 3000 parameter sets with faster PRL:PRLR:SOCS degradation values (**Figure 6A**, orange). We then simulated the model with the parameter sets from either group, comparing the predicted dynamics of the six quantities for which there is experimental data (**Figure 4**). We also considered one additional output, the SOCS protein, given its role in JAK-STAT signaling (see *Methods* section for more detail) and involvement in the receptor degradation rate. Interestingly, there was no clear difference between the predictions for the six quantities that have been measured experimentally due to sampling from the two groups of values for PRL:PRLR:SOCS degradation rate (**Supplementary Figure S9**).

**Figure 6.**
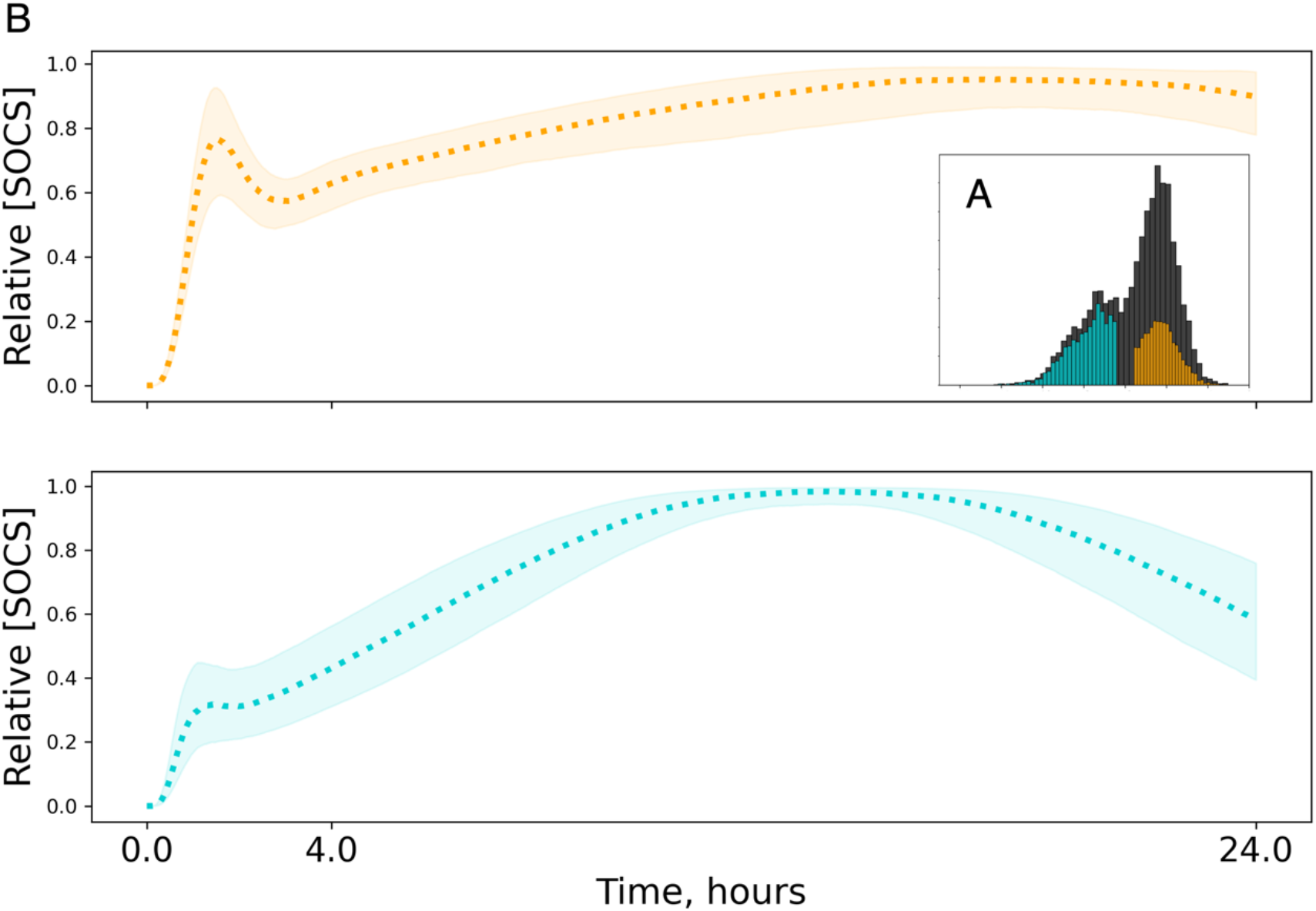
Predicted SOCS dynamics associated with the two parameter regimes for the rate of degradation of the PRL:PRLR:SOCS complex. (A) The posterior distribution indicates either a fast or slow PRL:PRLR:SOCS degradation rate. (B) Simulations produced by sampling from the two parameter regimes resulted in distinct SOCS dynamics in both the short term (0-4 hours) and the long term (up to 24 hours). Dashed line, median prediction; shaded region, inter-quartile range. Orange, predictions sampled from faster degradation regime; blue, predictions sampled from slower degradation regime.

However, two clearly distinct SOCS predictions emerged. The predicted SOCS dynamics associated with the slower receptor complex degradation exhibit little to no early peak and transient activation 24 hours after PRL stimulation (**Figure 6B**, blue). On the other hand, the SOCS predictions associated with faster receptor complex degradation exhibit a significant early peak and sustained activation 24 hours after PRL stimulation (**Figure 6B**, orange). Overall, we connected distinct SOCS signaling dynamics to the two plausible ranges of PRL:PRLR:SOCS degradation, thereby producing two hypotheses as to the dynamics of the SOCS protein.

Sampling from the posterior distribution produced a range of distinct predictions for the dynamics of the SOCS protein. Thus, we used the SOCS dynamics to characterize the model predictions. In addition to the two SOCS behaviors described above, the model predicts two more possible behaviors of the SOCS proteins – no early peak with sustained signaling and early peak with transient signaling. Thus, there are a total of four possible SOCS profiles. We used shape classification (see Methods) to determine the predicted frequency of each of these profiles. We applied this classification criterion to 10,000 predictions, generated by randomly sampling 10,000 parameter sets from the posterior. The resulting frequencies are displayed in the left panel of **Figure 7A**. An early peak in SOCS concentration followed by sustained activation emerges as the most probable SOCS profile; this behavior is illustrated in the right panel of **Figure 7A**. We next sought to test the validity of these hypothesized dynamics predicted by the model.

**Figure 7.**
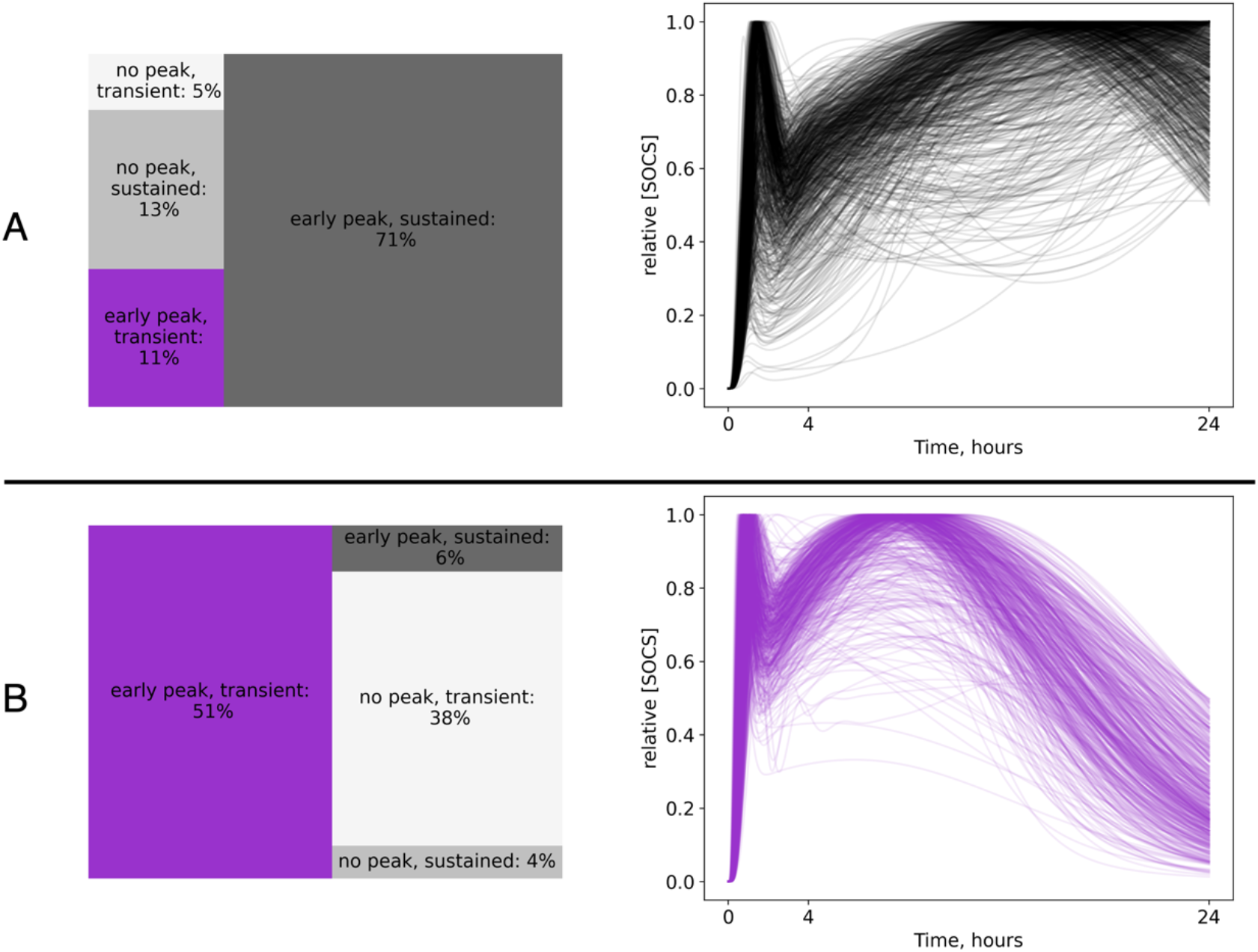
Comparison of the frequency of predicted SOCS dynamics. We determined how many predictions of SOCS dynamics are classified as no peak and transient activation (light gray), early peak and transient activation (dark gray), no peak and sustained activation (medium gray), and early peak and sustained activation (purple; target shape), before and after data integration (left). The most probable SOCS dynamics for each case are also displayed (right). (A) Posterior inferred with the baseline data; (B) Posterior inferred with the extended data.

#### 3.2.3 Refining Posterior Produces Better Predictions and Enables Hypothesis Testing

To confirm or reject our hypothesized SOCS profiles, we first conducted a literature search investigating SOCS dynamics in JAK2-STAT5 pancreatic beta cell signaling. Although there is significant interest in the topic, there exists no direct evidence of SOCS dynamics for our system of interest. However, our literature search did suggest that SOCS likely displays an early peak in its concentration followed by transient activation 24 hours post-stimulation with PRL (**Supplementary Figure S3**). We designated this as the target shape for the next phase of our investigation: model validation. More details on how we deduced this target shape are given in the Methods.

Conducting model validation, we found that our original model is inconsistent with the target shape. Our original model predicted sustained SOCS activation as the most probable behavior, while our validation metric indicates transient SOCS activation is more likely. Thus, we next we sought to establish a model that better captured the target SOCS behavior found in the literature. To do this, we aimed to better constrain the long-term SOCS dynamics, which requires experimental data at later timepoints. Although we could not find direct measurements of SOCS dynamics following PRL simulation in pancreatic beta cells, we were able to uncover new observations of the JAK2 and STAT5 proteins at later timepoints (personal communication with R.L. Sorenson). We incorporated these new data into our Bayesian inference procedure. We again ensured that (1) our posterior inference algorithm converged via convergence diagnostics (**Supplementary Figure S6B and S7B**) and (2) the updated model predictions recapitulated the training data (**Supplementary Figure S10**). We then validated our updated model against the target shape (**Figure 7B**).

The updated model predictions for SOCS are significantly different from the original predictions. Before integrating the new data for later timepoints, the most probable SOCS dynamics exhibit sustained activation with an early peak, and the literature-confirmed SOCS behavior of transient activation at 24 hours with an early peak accounts for only 11% of the simulations (**Figure 7A**, left). After data integration, more than 50% of the predicted SOCS dynamics exhibit the target SOCS behaviors—transient activation with an early peak (**Figure 7B**, left). Thus, the updated predictions agree with the target behavior better than the original predictions.

We find that when additional measurements for signaling proteins at later timepoints are used to inform the model parameters, different long-term SOCS dynamics are predicted (**Figure 8**). The percentage of the predicted long-term SOCS dynamics following PRL stimulation switched—from sustained (predicted with the original model) to transient activation (predicted with the updated model), as shown in **Figure 8A**. Specifically, transient activation accounts for 90% of predictions, where previously it accounted for only 15% of predictions. Based on this, as well as our literature-informed validation metric, we can be reasonably certain that SOCS displays transient activation in beta cell signaling.

**Figure 8.**
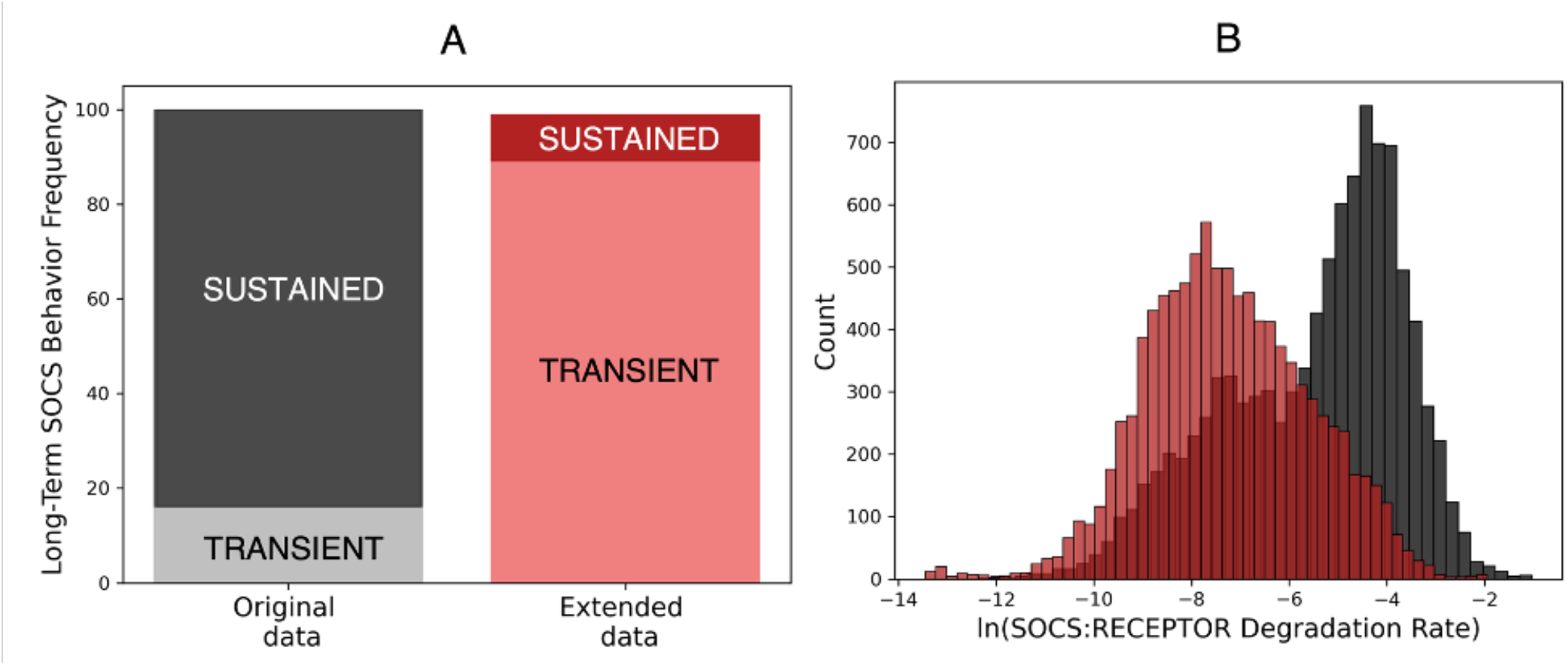
Comparison between the original and updated models’ posterior distribution and predictions. (A) Bar chart of long-term SOCS behavior classification frequencies before and after data integration. Black, original model (prior to data integration); red, updated model (after data integration). (B) Histograms of the marginal posterior distributions of the PRL:PRLR:SOCS degradation rate before and after integration of data at later time points, displayed on a natural-log scale.

Finally, the change in the SOCS predictions also corresponded with a significant change in the marginal posterior of the PRL:PRLR:SOCS complex degradation rate (**Figure 8B**), as well as smaller changes in the marginal posteriors of other parameters (**Supplementary Figure S11**). In the case of the PRL:PRLR:SOCS complex degradation rate, one mode of the distribution is eliminated after data integration. The mode remaining corresponds to the slower PRL:PRLR:SOCS degradation rate (colored blue in **Figure 6**). Overall, our updated model predicts transient SOCS activation and favors a slower PRL:PRLR:SOCS degradation rate.

## 4. Discussion

Our work establishes an improved approach to Bayesian hypotheses formation, refines an existing model of pancreatic beta cell signaling, and generates novel biological insights. We introduced a KL divergence-based ranking, an approach that facilitates hypothesis generation by prioritizing parameters for subsequent investigation. While other approaches for hypothesis formation using posteriors exist, they are largely qualitative and can be time consuming, as they typically require qualitative evaluation of every model parameter (Hug et al., 2013; Thijssen et al., 2018; Tötsch and Hoffmann, 2020). Our approach streamlines the qualitative investigation of model parameters by providing a means to systematically search the posterior distribution for different kinds of alternative hypothesis evidence We prioritize this subset of parameters by how much information they gain from the data using KL divergences and further group the parameters using K-means clustering. For larger models, K-clustering the divergences is especially useful in selecting only a subset of parameters for analysis, thereby further expediting subsequent investigations. Prioritizing parameter estimates that gain significant information from the data allows us to focus on new information, which might contain evidence of alternative model hypotheses. Importantly, our approach is versatile. For example, K-means clustering is an unsupervised learning technique (Aldenderfer and Blashfield, 1984; Bible et al., 2013). Our approach is therefore applicable to models with KL divergence patterns unknown *a priori*. In addition, KL divergence may be estimated between marginal prior and posterior distributions of any form (Kullback and Leibler, 1951). We provide examples of KL divergence estimated using both a log uniform and log normal prior distribution; however, our approach is also applicable to models with prior and posterior formulations different than those used here. Finally, KL divergence offers a means to measure the information gained from new data, without knowing *a priori* how this information will be reflected in the posterior. This means that it may be used to detect different manifestations of alternative hypothesis evidence, such as a bimodal posterior or a posterior informed in a contradictory manner by different datasets.

We tested our approach with two examples, which together showcase our approach’s utility in systematically uncovering different types of alternative hypothesis evidence. First, we apply our KL divergence ranking to an established example of Bayesian hypothesis formation previously presented in the literature (Tötsch and Hoffmann, 2020). Then, we apply our ranking in a novel study of a computational model of prolactin-induced JAK2-STAT5 signaling, which was previously established in in Mortlock et al. (Mortlock et al., 2021) and most recently used to investigate cell heterogeneity (Simoni et al., 2022). In both examples, we used KL divergence to rank parameters by how much information they gain from experimental data. The purpose of this ranking is to guide subsequent model investigations: rather than searching all model parameters at random, it may be used to prioritize searching the parameters that gained the most information from the data for evidence of alternative model hypotheses. In the first example, our approach ranks first the target parameter, which is the same parameter whose posterior was previously found to be inconsistent between data sets. In the second example, we cluster our KL divergence ranking to select only a subset of parameters to search for qualitative evidence of alternative hypotheses, decreasing the number of parameters we search from 33 to 3. Within this subset, we find a bimodal posterior, indicating two plausible regimes for the rate of degradation of the SOCS-bound receptor. These regimes characterized unique SOCS dynamics, which formed testable hypotheses of beta cell signaling mechanisms. Overall, these results demonstrate our approach’s utility in systematically uncovering different types of alternative hypothesis evidence, enabling the generation of mechanistic hypotheses explaining complex protein signaling networks. Similar studies concluded after proposing such alternative hypotheses (Hug et al., 2013; Tötsch and Hoffmann, 2020). In this work, we moved beyond hypothesis formation by testing *in silico* the proposed SOCS dynamics.

While there has been significant research investigating SOCS’ role in the canonical JAK-STAT pathway, as well as the role of SOCS in PRL-mediated JAK2-STAT5 signaling, no observations of SOCS dynamics for our exact system of study exist (Chong et al., 2001; Croker et al., 2008; Jiao et al., 2013; Millette et al., 2022; Ram and Waxman, 1999; Rieck et al., 2009; Rønn et al., 2002; Ye and Driver, 2016). Thus, determining the dynamics of SOCS in response to PRL is an open challenge in pancreatic beta cell biology. To investigate this question, we refined our posterior by incorporating additional data at extended timepoints. We then validated our updated model by ensuring its predictions were consistent with dynamics of the CISH and SOCS2, which are shown to be conserved across experimental conditions (Chong et al., 2001; Croker et al., 2008; Ram and Waxman, 1999). In contrast, the model obtained prior to integrating the new experimental data was not consistent with these target dynamics. This result established the predictive capabilities of our updated model. The model predicts that SOCS likely displays transient activation after PRL stimulation. Because SOCS provides negative feedback to the JAK-STAT pathway, it plays a crucial role in controlling hormone-mediated beta cell proliferation. Thus, this insight is useful for developing strategies to enhance JAK2-STAT5 signaling and, ultimately, a therapeutic target for Diabetes treatment: beta cell proliferation (Durham et al., 2019).

In addition to determining the most probable SOCS dynamics in response to PRL stimulation, our analysis of the posterior distribution revealed insight about the degradation rate of the SOCS-bound receptor. We investigated how the new data changed the probability of different SOCS-bound receptor degradation rates by comparing the marginal distribution of the SOCS-bound receptor degradation rate before and after data integration. After data integration, the probability of a faster degradation rate decreased, and the slower degradation regime emerged as most plausible. Overall, our analysis of the posterior before and after data integration complements how it has been used in previous works. For example, this simple approach has been used to understand the information in different types of data and to demonstrate model inadequacy (Thijssen et al., 2018; Tötsch and Hoffmann, 2020).

We acknowledge some limitations and future improvements of our KL divergence based approach as well as our work more generally. First, while our KL divergence-based search out-performed a multi-modality search in the JAK-STAT case study, this may not always be the case, depending on the specific combinations of prior and posterior distributions. Our KL divergence ranking is sensitive to cases in which the prior and posterior have significant sections of non-overlapping mass. However, there are cases in which alternative hypothesis evidence may not manifest in this manner. For example, if the bimodal SOCS-bound receptor degradation rate posterior was shifted leftwards, such that it was centered under the prior distribution, the KL divergence from the prior and posterior would be reduced, perhaps leading to this parameter being excluded from the cluster that was most influenced by the data. In this case, posterior analysis could be prioritized based solely on KL divergence ranking, as we showed in the Totsch test case. However, it may be just as efficient to search the 12 parameters identified by the multi-modality search. Despite this kind of exception, our approach does provide key advantages over a multi-modality search. For example, our approach searches for a more general class of alternative hypothesis evidence, including both bimodalities and posteriors informed in a contradictory manner by the different data sets. This is particularly useful if the form of evidence is not known *a priori*. In addition, in the case that a posterior has a significant mode outside the prior, our method will highlight this bimodality above the rest. This is useful in determining which bimodality, perhaps of many, to investigate further. A second limitation of our approach is that while the clustering results we present here are biologically meaningful, this may not always be the case. Clustering analysis seeks to identify latent structures in data. However, one must consider whether the structure it identifies is “real” or simply imposed on the data by the method (Aldenderfer and Blashfield, 1984; Bible et al., 2013). Here, our top data-informed cluster was deemed meaningful as its members shared a biological mechanism related to receptor degradation. Future work should similarly assess the meaning of clustering results. If no meaningful clusters emerge, one could instead prioritize posterior analysis based solely on KL divergence ranking, as we did in our first test case.

A limitation of our work more generally relates to how well the JAK-STAT model captured experimental data. While our JAK-STAT model calibrated with either the original data or with the extended data recapitulate most of the experimental data, a systematic deviation between model predictions and the experimental data does exist at the 4-hour timepoint. This systematic deviation indicates that our model’s predictive accuracy for the 4-hour timepoint is low. This is less important for our investigation, as we were primarily concerned with the models’ predictive accuracy prior to the 4-hour timepoint (representing the short-term signaling response) and at the 24-hour timepoint (reflecting the long-term response that influences cellular behavior). Further, in the context of our investigation, these models are useful not only for their predictive accuracy, but also their ability to generate, test, and refine hypotheses (Enderling and Wolkenhauer, 2021). However, future investigations could consider changes to the model structure to better recapitulate the 4-hour timepoint. Another limitation of our work relates to shape classification of the species’ timecourses. While we based our SOCS shape classification criterion on several western blot data sets, there remains an element of ‘modelers choice’ when defining these criteria. If our results were highly sensitive to the criterion definition, this would limit their importance. To address this limitation, we varied both of our classification criteria and ensured our results were conserved. This result also verifies that the transition in the SOCS’ response as the PRL:PRLR:SOCS degradation rate varies is well-described by a discrete classification. A third limitation is that while we conducted a comprehensive literature search and developed a validation for our model with inferable literature evidence, this evidence is less certain than direct observations of SOCS. Future work could focus on collecting western blot measurements of SOCS, particularly from 0-4 hours, to determine whether SOCS displays an early peak in activation. Lastly, although rodent models provide valuable insights into human biology, there are certainly limits to the applicability of these insights(Baeyens et al., 2016; Salazar-Petres and Sferruzzi-Perri, 2022). Our mathematical model is calibrated with data collected in the INS-1 rat cell line (Brelje et al., 2004, 2002; Fujinaka et al., 2007). Though the net effects of pregnancy on beta cells, the expansion of beta cell mass and increased insulin secretory capacity are conserved between rodent and human beta cells, it is unclear how the mechanisms driving these changes in rodent cells map to the mechanisms driving these changes in human cells. Future work can test the applicability of our insights in human beta cell biology.

## 5. Conclusions

Overall, our work establishes an improved approach to hypothesis formation using the Bayesian posterior, refines an existing model of pancreatic beta cell signaling, and ultimately answers an open question in the field of pancreatic beta cell biology. Our approach to search for alternative hypothesis evidence provides a basis for other investigations of chemical reaction networks, including other biochemical signaling networks. The refined model provides a quantitative framework for future investigations of beta cell signaling. Finally, the novel insights into pancreatic beta cell biology generated here can potentially be leveraged to develop strategies to increase functional beta cell mass.

## Acknowledgements

The authors acknowledge members of the Finley research group for constructive feedback and both Niklas Totsch and Robert Sorenson for their generous help in accessing the data presented in their respective papers.

## CRediT authorship contribution statement

**Holly A. Huber:** Conceptualization, Formal analysis, Investigation, Methodology, Writing – original draft, Writing – review & editing. **Senta K. Georgia:** Writing – review & editing. **Stacey D. Finley:** Conceptualization, Supervision, Resources, Writing – review & editing.

