## Supplementary Materials for "Systematic Bayesian Posterior Analysis Guided by Kullback-Leibler Divergence Facilitates Hypothesis Formation"

### 1. Additional Details Regarding Kullback Leibler Divergence Approximation

We estimate the Kullback-Leibler (KL) Divergence from the marginal prior distribution to the marginal posterior distribution for each inferred parameter given the original data. KL Divergence provides an idea of the difference between distributions—thus, the KL Divergence from the marginal prior distribution to the marginal posterior distribution provides an idea of how different our inferred posterior is from our original prior distribution.

More formally, KL divergence is defined:

$$D_{KL}(P|Q) = \int_{-\infty}^{\infty} p(x) \log\left(\frac{p(x)}{q(x)}\right) dx$$

*Eq. ( S.1 )*

In words, the KL Divergence from Q to P is the expectation of the logarithmic difference between the probabilities P and Q, where the expectation is taken with respect to P (Kullback and Leibler, 1951).

We leverage two approximation techniques to estimate the KL divergence. We begin with the analytical form of our prior distribution and samples, generated from our Bayesian inference procedure, from the posterior distribution. To calculate the KL divergence, we need to estimate the probability density function of the marginal posterior distribution using these samples. Kernel density estimation provides such an estimate. After log-transformation, samples from our marginal posterior distributions are approximately normal (see figure above). Thus, we choose to use a gaussian kernel parameterized with Silverman's bandwidth in our kernel density estimate. Representative figures shown below provide a comparison between the samples drawn from our kernel density estimates and the samples from the marginal posterior distributions. Because these two sets of samples are essentially identical for each marginal posterior distribution, we conclude our kernel density estimate provides an acceptable approximation of each marginal posterior distribution's probability density function.

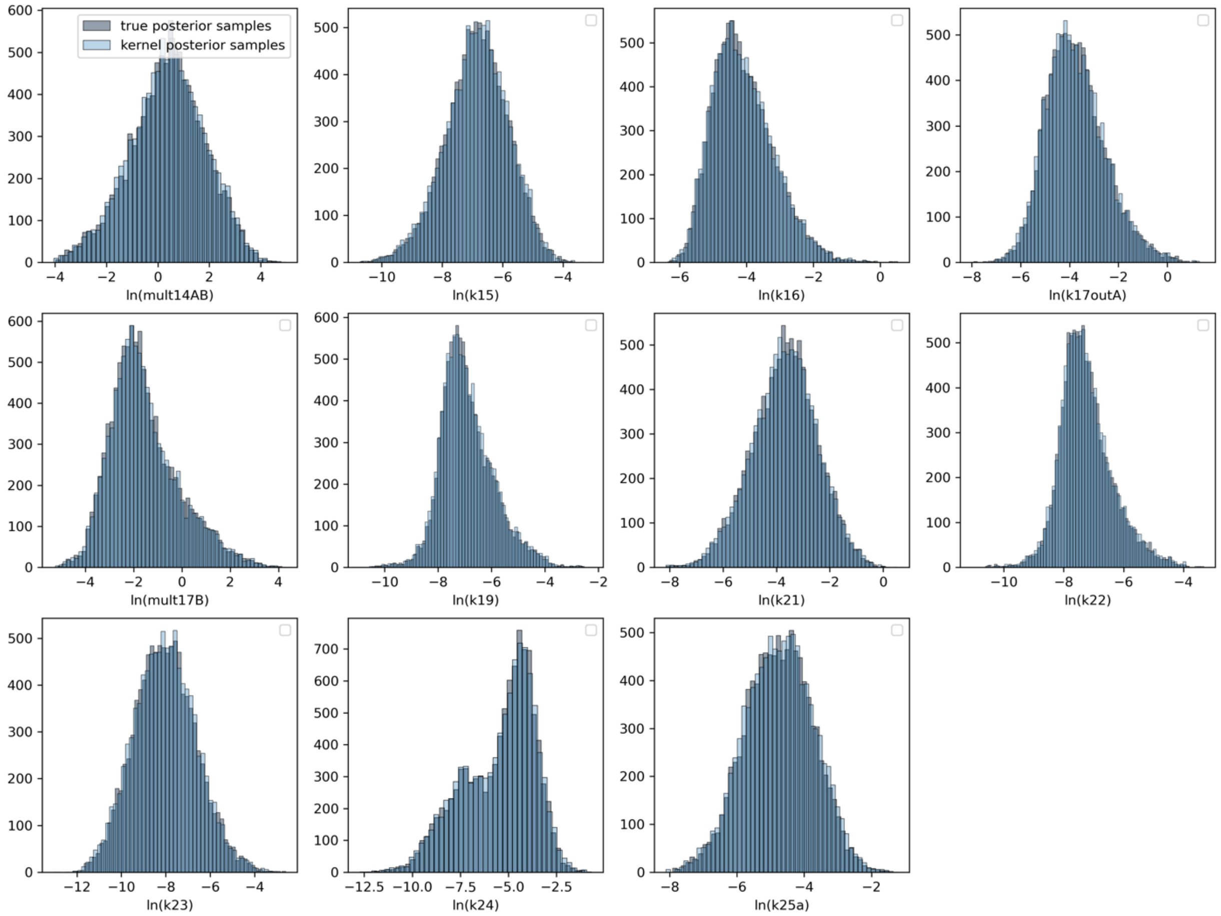


**A representative sample of histograms assessing accuracy of kernel density estimates.** Light blue; samples generated from our kernel density estimate of each log-transformed, marginal posterior distribution; grey, samples from the log-transformed, marginal posterior distribution.

The second approximation technique we use is a simple application of the Law of Large Numbers. The Law of Large Numbers states that the sample mean approaches the true expected value as the number of samples approaches infinity. Because the KL Divergence is an expectation, we may thus approximate it with a sample mean, calculated with samples from the posterior distribution:

$$D_{KL}(P|Q) = \int_{-\infty}^{\infty} p(x) \log\left(\frac{p(x)}{q(x)}\right) dx = \frac{1}{n}\sum_{i=1}^{n} \log\left(\frac{p(x_i)}{q(x_i)}\right)$$

*Eq. ( S.2 )*

### 2. Additional Details Regarding Posterior Inference

#### *2.1 Background on Bayesian Inference*

Bayesian inference is a distribution approximation problem in which we are trying to approximate the conditional probability of the model parameters given experimental data, ie. the posterior distribution. The posterior distribution is proportional to the likelihood distribution multiplied by the prior distribution. This relationship is specified via Bayes Theorem:

$$P(\vec{\theta}|\vec{x}) \propto P(\vec{x}|\vec{\theta}) * P(\vec{\theta})$$

*Eq. ( S.3 )*

that is,

$$posterior \propto likelihood * prior$$

where:

$$\vec{\theta} = vector\ of\ model\ parameters$$
$$\vec{x} = vector\ of\ experimental\ data$$

#### *2.2 Approximation Implementation Details.*

To generate samples of our posterior distribution (here, a posterior sample is a vector of the 33 inferred kinetic rate constants), we implemented the Metropolis-Hastings algorithm. This algorithm generates an aperiodic and irreducible Markov chain that satisfies the detailed balance condition. The Markov chain generated by the Metropolis-Hastings algorithm is aperiodic as the algorithm always allows for sample rejection. To create an irreducible Markov chain, we chose to use a lognormal proposal distribution. Because the support of the lognormal distribution (all positive real numbers) includes the support of the posterior, we ensure irreducibility. By generating an aperiodic and irreducible Markov chain, we guarantee asymptotic convergence to the posterior distribution (Andrieu et al., 2003). To ensure convergence in practice, we use established quantitative (split-Gelman-Rubin diagnostic) and qualitative (trace plots, autocorrelation plots, and histograms) metrics (Cowles and Carlin, 1996; Gelman, 2014).

We approximate two posteriors: the distribution of parameters given the original data and the distribution of parameters given the extended data. **Supplementary Figure S10** visualizes and compares our marginal prior distributions and marginal posterior distributions for each data set. In both cases, we ended sampling after meeting our convergence threshold. In the case of the original posterior, we initialize five independent Markov chains from different initial conditions. Each chain generated ~300,000 samples prior to convergence; thus, in total, we used ~1,500,000 samples to approximate the posterior given the original data. In the case of the refined posterior, we initialized three independent Markov chains from different initial conditions. Each chain generated ~200,000 samples prior to convergence; thus, in total we used ~600,000 samples to approximation the posterior given the extended data.

***2.3 Details regarding convergence criteria.***

The Gelman-Rubin (G-R) method monitors convergence by comparing the variance between difference sample chains and the variance of the pooled sample chains (Brooks and Gelman, 1998). Chains are initialized at different locations of the posterior; thus, the variance between sample chains is larger than the variance of the pooled chains. However, as the chains sample the same posterior distribution, the variance between sample chains approaches the variance of the pooled sample chains. Quantitatively, this means that as the chains converge, the ratio of these variances (the G-R metric) approaches one.

For the G-R method to be applicable, two key assumptions must be met. First, the chains must be initialized from a distribution over-dispersed with respect to the posterior distribution. To do this for our original posterior estimation, we first initialized our chains from the prior distribution used in Mortlock et al. This prior distribution is derived from an aggregated dataset of $k_{cat}$ parameters across cell types and intracellular cell pathways (Bar-Even et al., 2011).Thus, because this model's posterior distribution describes one cell type and one pathway, this distribution was assumed to be over-dispersed with respect to our posterior distribution. Because we are particularly interested in the SOCS-bound receptor degradation rate, we also initialized chains from either extreme for this parameter. To initialize overdispersed chains for the refined posterior estimation, we initialized chains from either extreme of the SOCS-bound receptor degradation rate. This is because we were particularly interested in the change of this degradation rate after incorporating new data into our estimate.

The G-R method also assumes that convergence in the first and second moments (mean and variance) implies convergence in the distribution. Gaussian distributions are completely defined by their first and second moment, thus, the G-R method is applicable for Gaussian or approximately Gaussian distributions (Brooks and Gelman, 1998). After applying a log-transformation of our marginal posterior distributions estimated with both the original data and extended data, we utilized Q-Q plots to verify that the distributions were approximately Gaussian (representative plots are shown below). To do this, we checked that the sample quantiles match the theoretical quantiles for 99% of the data, ie. between z-scores from -3 to 3. By this assessment, most of the distributions may be considered approximately Gaussian, and therefore the G-R method is an applicable metric of convergence for most of our parameters.

Though the majority of our marginal posterior distributions are approximately Gaussian, there were a few distributions that showed signs of being non-Gaussian. For example, the Q-Q plot of the parameter mult17B indicates skewness while the Q-Q plot of bimodal parameter k24 indicates a thin tail. Thus, we took additional steps to ensure that the G-R diagnostic would provide a suitable metric of algorithm convergence for our posterior. First, we used the *split-R* G-R diagnostic for all inference protocols. The split-R diagnostic improves upon the traditional G-R diagnostic by comparing the first half of each chain to the second half, thereby allowing the detection of within-chain non-convergence (Gelman, 2014; Vehtari et al., 2019). Second, we initialized additional chains, for a total of five chains, for the inference protocol given the original data. That is, when we assessed the convergence of a multimodal distribution. In a study comparing convergence diagnostics, the G-R diagnostic outperformed other metrics in diagnosing non-convergence for multi-modal posteriors (Cowles and Carlin, 1996). This was because of the multiple chains used to assess convergence. By increasing the number of sampling chains, we increase the likelihood of uncovering nonconvergence for multimodal posteriors, like k24. With these additional steps, the G-R diagnostic also provides an applicable metric of convergence for the non-Gaussian posteriors of our Bayesian inference protocol.

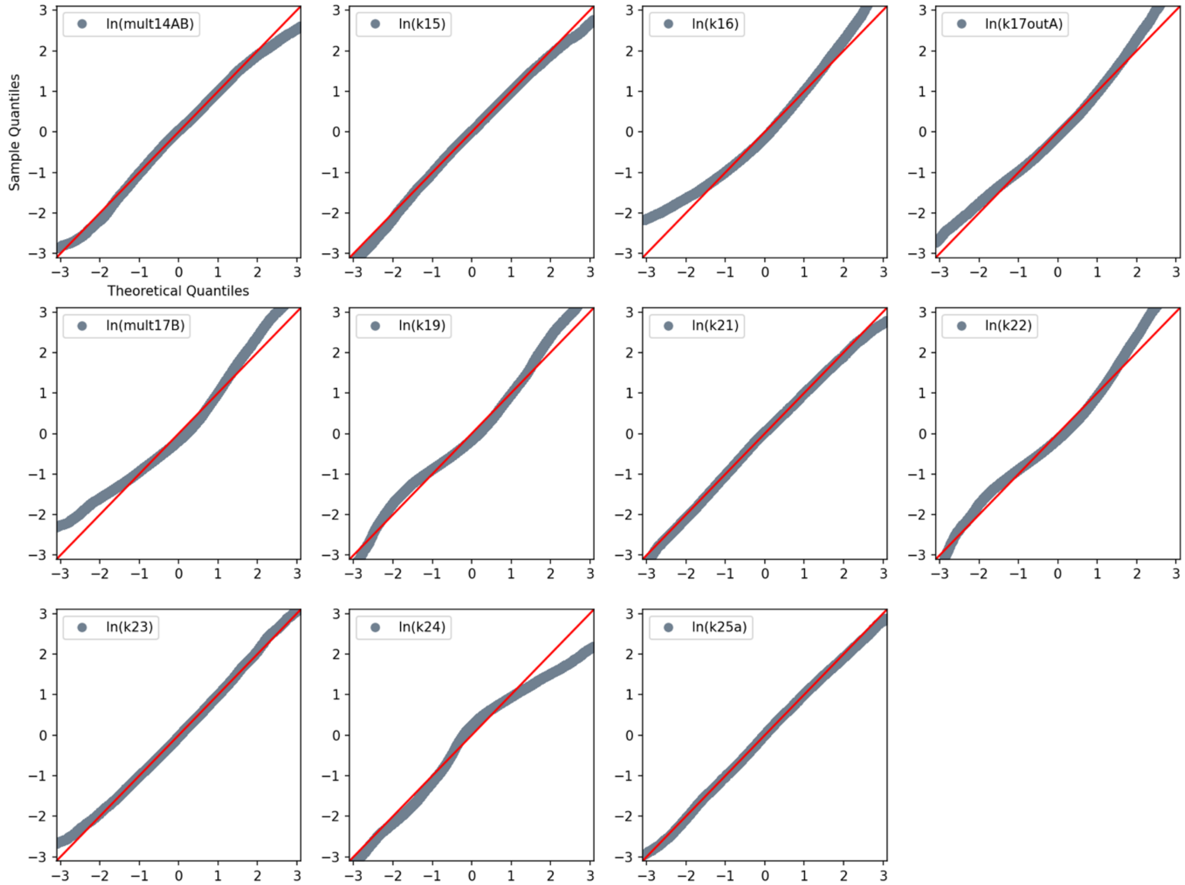


**A representative sample of Normal quantile-quantile (Q-Q) plots.** Plots assess the normality of log-transformed marginal posterior distributions. Samples from each marginal posterior distribution are plotted against the theoretical quantile of a normal distribution parameterized by the marginal posterior distribution's sample mean and sample variance. Grey dots, samples from a marginal posterior distribution versus its theoretical quantiles; red line, theoretical quantiles versus theoretical quantiles.

### 3. Additional Details Regarding K-Means Clustering

We use K-means clustering to group our $n$ = 33 KL divergences into $k$ disjoint clusters, which are described by the mean $\mu_j$ of the samples in the cluster. The objective of this assignment is to minimize the sum of the squared distances between each cluster's mean and member data points—i.e., the within-cluster sum of squares criterion (WCSS):

$$\sum_{i=1}^{n} \min_{\mu_j} \left( \left\| x_i - \mu_j \right\|^2 \right)$$

*Eq. ( S.4 )*

To implement K-means clustering, one must specify the number, *k*, of clusters. We allowed *k* to range from 1 to 7. The optimal number of clusters was set based on two diagnostics: a qualitative diagnostic, the ‘Elbow Method’, and a quantitative metric, the average Silhouette Coefficient (Jolly, 2018; Rousseeuw, 1987). To create the Elbow diagnostic, we plotted the WCSS against the number of clusters used for a K-means fit. At first, the WCSS decreases dramatically with increasing cluster numbers; however, eventually the WCSS starts decreasing only marginally with increasing cluster numbers (**Supplementary Figure S2**). The ideal number of clusters occurs when the trend in the WCSS switches. Graphically, this point looks like an elbow.

To calculate the Silhouette Coefficient, we used scikit learn’s *silhouette_score* function (Pedregosa et al., 2011). The Silhouette Coefficient is defined for each sample—here a KL divergence value. Each coefficient is bounded from -1 to 1 and a higher coefficient indicates dense and well-separated clusters (Rousseeuw, 1987). We take the average Coefficient over all 33 samples; the ideal number of clusters occurs at the highest Silhouette Coefficient. Finally, we note that the Silhouette Coefficient always equals 0 when k=1; thus, it cannot determine whether *k* = 2 or *k* = 1 better describes the data. However, the Elbow Method does not have this limitation. Thus, by using both diagnostics to determine the optimal number of clusters, we can be confident in our determination.

### 4. Supplementary Tables and Figures

**Table S1.** Parameters defining likelihood, prior, and proposal distributions. See below for why we implemented a natural log-transformation for the means of the lognormal distributions.

<table>
<tr><th>Likelihood of Experimental Data Point, j</th><th>Likelihood of Experimental Data Vector</th></tr>
<tr><td>$x_j|\vec{\theta} \sim \mathcal{N}\left(\mathcal{M}(\vec{\theta})_j, \sigma^2\right)$</td><td>$\vec{x}|\vec{\theta} \sim \prod_j \mathcal{N}\left(\mathcal{M}(\vec{\theta})_j, \sigma^2\right)$</td></tr>
<tr><td colspan="2">$\sigma^2 \sim Inverse\ Gamma(2, 0.001)$<br>$x_j = experimental\ data\ point\ j$<br>$\vec{\theta} = vector\ of\ model\ parameters$<br>$\vec{x} = vector\ of\ experimental\ data$<br>$\mathcal{M}(\vec{\theta})_j = ODE\ model\ prediction\ for\ data\ point\ j, evaluated\ with\ parameter\ vector\ \vec{\theta}$</td></tr>
<tr><th>Prior Distribution of Parameter, i</th><th>Prior Distribution of Parameter Vector</th></tr>
<tr><td>$\theta_i \sim LogNormal(\ln(\theta_{i0}), 2)$</td><td>$\vec{\theta} \sim \prod_i LogNormal(\ln(\theta_{i0}), 2)$</td></tr>
<tr><td colspan="2">$\theta_i = parameter\ i$<br>$\theta_{i0} = initial\ guess\ for\ parameter\ i$</td></tr>
<tr><th>Proposal Distribution of Parameter, i**</th><th>Proposal of Parameter Vector</th></tr>
<tr><td>$\theta_{i,n}|\theta_{i,n-1} \sim LogNormal(\ln(\theta_{i,n-1}), 0.1)$</td><td>$\vec{\theta}_n|\vec{\theta}_{n-1} \sim \prod_i LogNormal(\ln(\theta_{i,n-1}), 0.1)$</td></tr>
<tr><td colspan="2">$\theta_{i,n-1} = \ posterior\ sample\ of\ parameter\ i\ \ from\ Metropolis - Hastings\ iteration\ n - 1$</td></tr>
</table>

Derivation S1. Deriving natural log-transformed means for parameterizing lognormal distributions:

Our formulation purposefully used the log-parameters to parameterize the lognormal distribution. The following derivation details why.

First, we establish these definitions of the lognormal distribution:

By definition, the logarithm of samples generated by the lognormal distribution are normally distributed, with mean, $\mu$, and standard deviation, $\sigma$.

The median of this lognormal distribution is:

$$(1)\quad \text{median} = \exp(\mu)$$

Next, we establish how the *np.random.lognormal* function, which we use here to generate our pdf, is parameterized:

The *np.random.lognormal* function takes two inputs, the aforementioned $\mu$ and $\sigma$.

Finally, we establish our desired lognormal distribution shape:

We want a lognormal distribution centered around our initial parameter guess, $\theta_{i0}$

As explained in the manuscript, we set $\sigma$=2. What is left is to derive the parameterization of *np.random.lognormal* in terms of $\mu$ such that our desired log-normal distribution is generated:

To approximately center our lognormal distribution on $\theta_{i0}$, we set the median of the lognormal distribution equal to $\theta_{i0}$:

$$(2)\quad \text{median} = \theta_{i0}:$$

Plugging in our definition for the median of the lognormal distribution, (1):

$$(3)\quad \exp(\mu) = \theta_{i0}$$

Solving for $\mu$:

$$(4)\quad \mu = \ln(\theta_{i0}).$$

Thus, to correctly parameterize the *np.random.lognormal* function such that the resulting distribution is approximately centered at $\theta_{i0}$, we set $\mu = \ln(\theta_{i0})$.

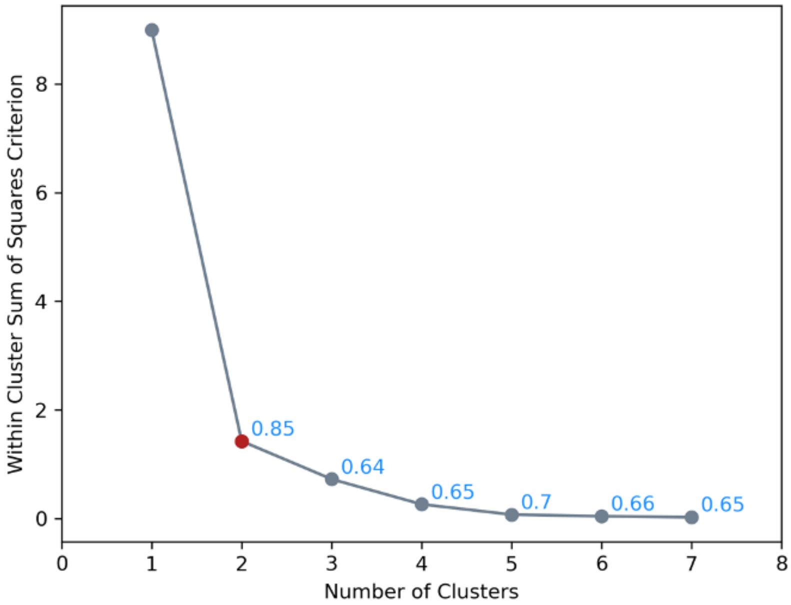


**Figure S2. Elbow Plot and Average Silhouette Scores for K-means clustering of KL divergences.** The Within Cluster Sum of Squares Criterion (WCSS) and average Silhouette Score (SS) was calculated for seven fits of the K-means algorithm to the KL divergence rankings. Each of these seven fits was initialized with a different number of clusters, k = {1,2,3,4,5,6,7}. Note that the average SS is not applicable for k = 1, thus, it is excluded for this plot. Grey line, WCSS for the fitted clusters; red dot, 'elbow' indicating the number of clusters that best explains the data; blue label, SS for fitted cluster.

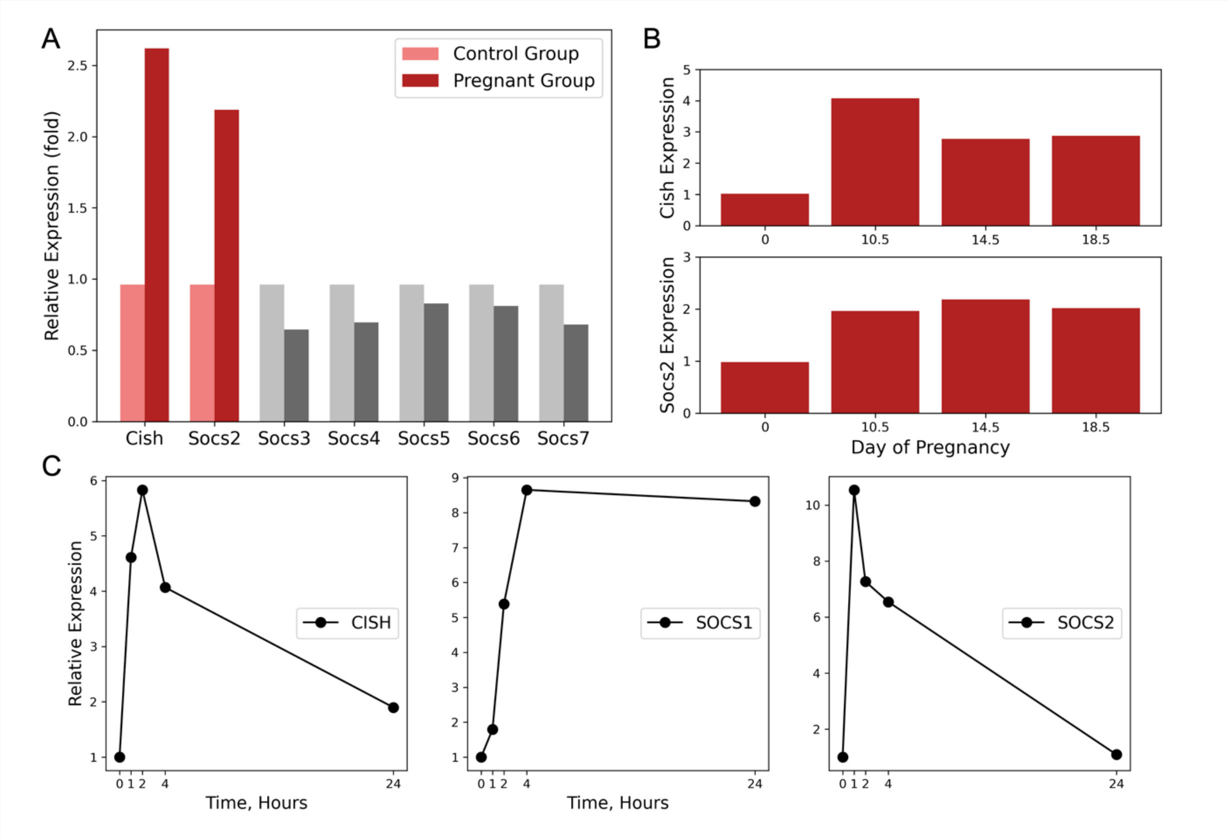


**Figure S3**. **Literature evidence of SOCS dynamics in pancreatic beta cells.** (A) mRNA expression for SOCS family of protein in pregnant and non-pregnant mice. Measurements were collected day 14.5 of gestation. Measurements are from Jiao (2013). (B) mRNA expression of Cish and Socs2 at day 0, 10.5, 14.5, and 18.5 of gestation. Measurements are from Rieck (2009). (C) Relative protein expression of CISH, SOCS1, and SOCS2 in pancreatic beta cells after stimulation with IN-Fγ. Measurements are from Chong (2001). Dark red and dark grey, pancreatic beta cells from pregnant mice; light red and light grey, pancreatic beta cells from non-pregnant mice; black, relative CISH, SOCS1, and SOCS2 concentrations

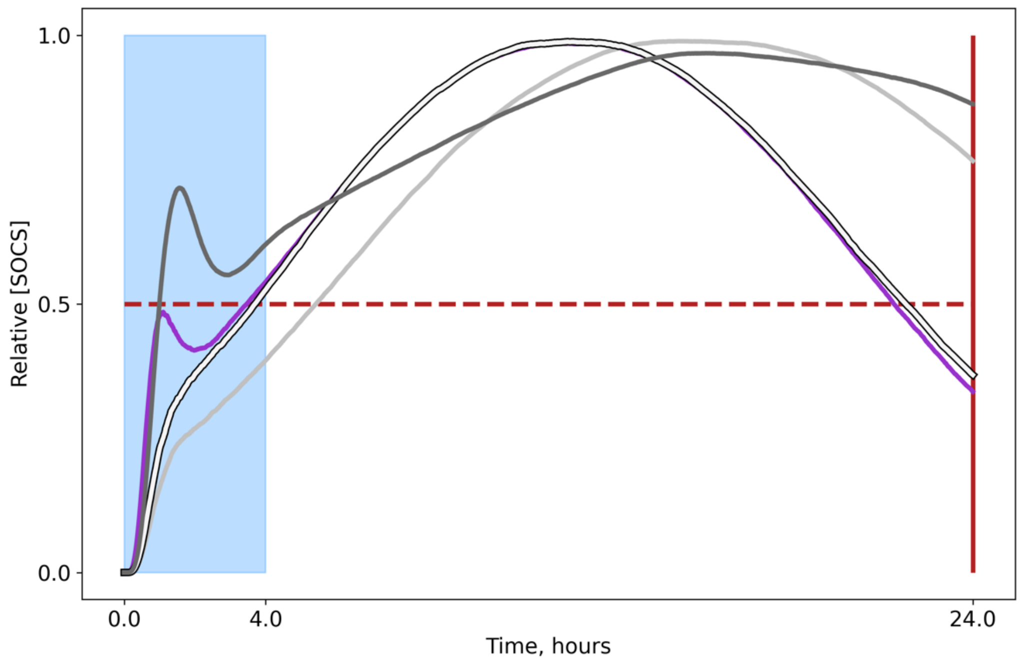


**Figure S4. Shape classification criteria applied to predicted SOCS time courses.** Predictions are categorized as having a peak in activation if a peak is present from 0-4 hours, otherwise, they are categorized as no peak. Predictions are categorized as sustained if SOCS levels are above 50% of max activation at 24 hours, otherwise, they are categorized as transient. Blue, short-term behavior time window; Red solid line, time at which long-term behavior is assessed; red dashed line, long term behavior threshold. Purple, early peak and transient; dark grey, early peak and sustained; light grey, no peak and sustained; white, no peak and transient.

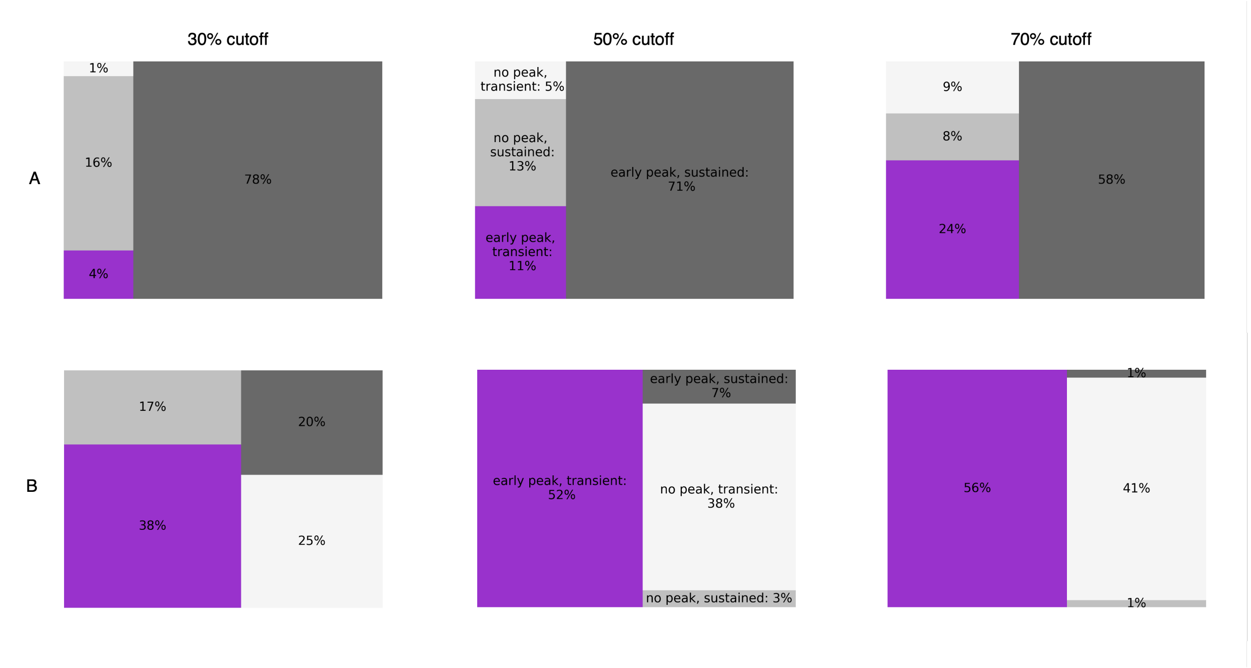


**Figure S5**. **Shape classification robustness.** Demonstration of results' robustness to changing shape classification criterion. Changing the early peak search window to 0-3 hours from 0-4 hours does not change the shape classification percentages shown in column 2. We also consider that the SOCS level at 24 hours must be less than 30%, 50%, or 70% of the maximum to be considered transient activation. (A) model trained with original data (B) model trained with extended data. Purple, early peak and transient; dark grey, early peak and sustained; light grey, no peak and sustained; white, no peak and transient.

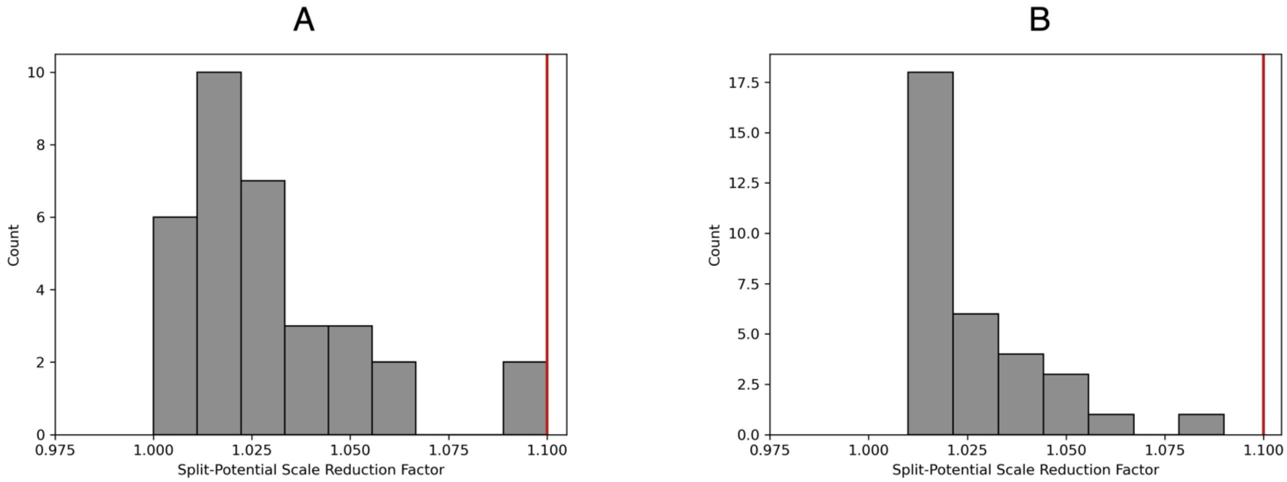


**Figure S6**. **Histogram depicts calculated $split - \hat{R}$ values for all 33 inferred model parameters.** Convergence was concluded when $split - \hat{R} < 1.1$ for all marginal posterior distributions. (A) baseline data, (B) extended data. Grey bars; $split - \hat{R}$ values for all 33 inferred model parameters. Red line; convergence threshold.

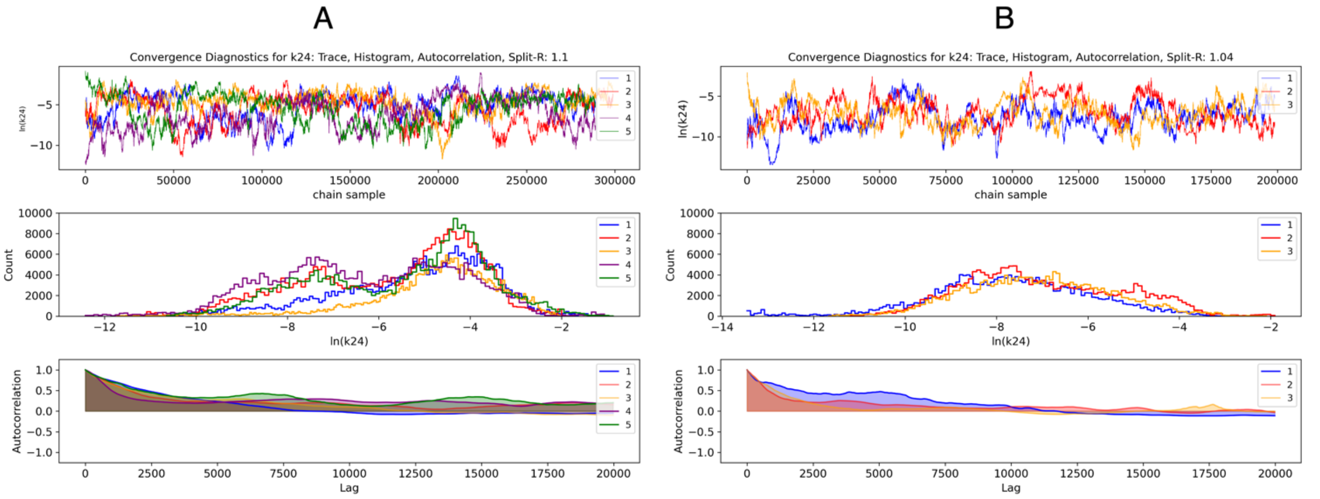


**Figure S7. Qualitative convergence diagnostic plots for the k24 marginal posterior distribution**. (A) baseline data, (B) extended data. Three independent Markov chains are indicated in blue, yellow, and red. $Split - \hat{R}$ diagnostic is included in the figure header. From top to bottom, plots include (1) chain traces vs sample iteration (2) histogram of chain samples (3) chain autocorrelation vs. lag and (4) chain summary statistics.

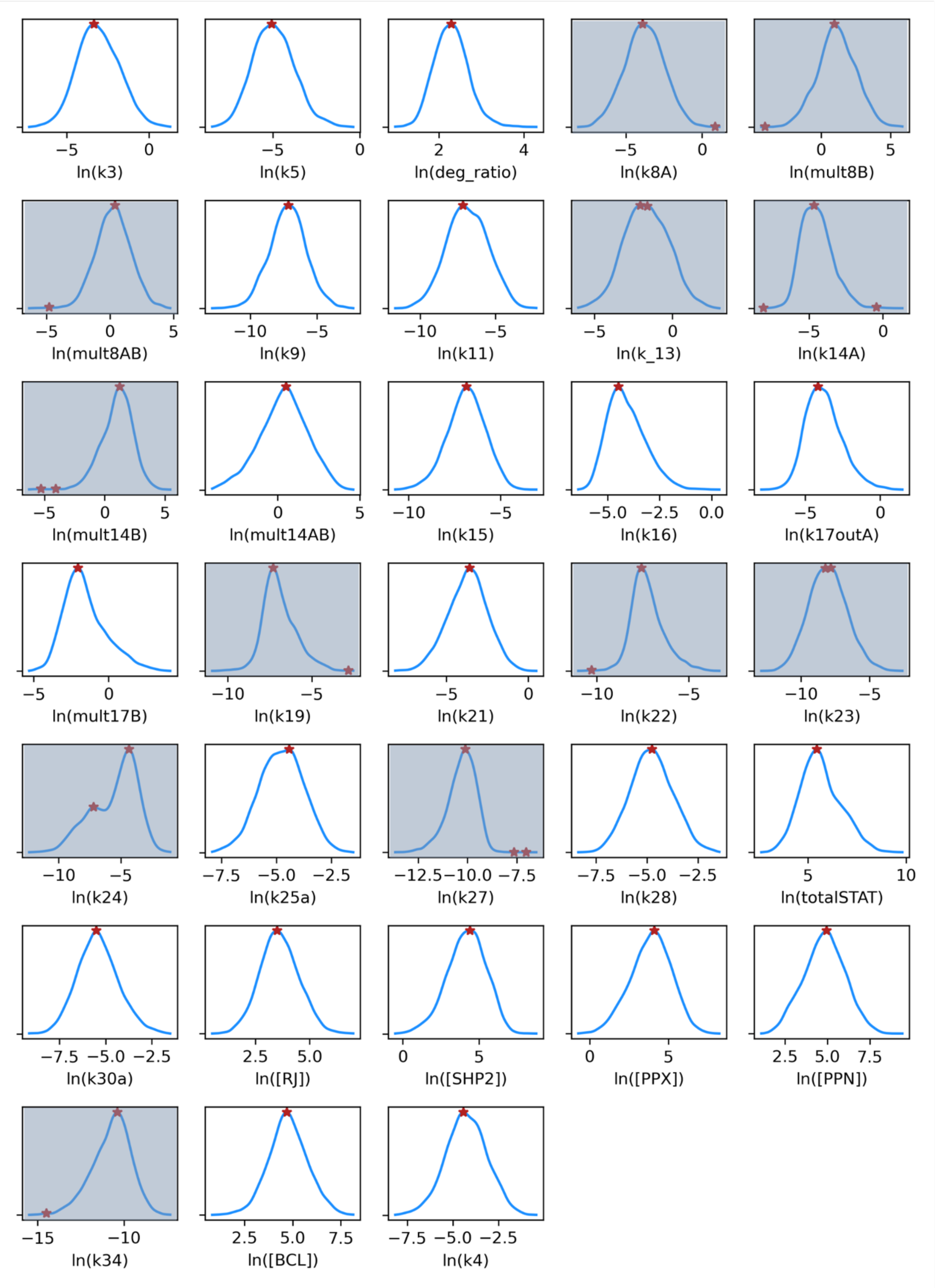


**Figure S8. Multimodality Search**. Results of multi-modality search for all 33 model parameters. Detected modes are noted in red, KDEs used for finding peaks are plotted in blue. In total, 12 modes are detected; those parameters are indicated by gray background.

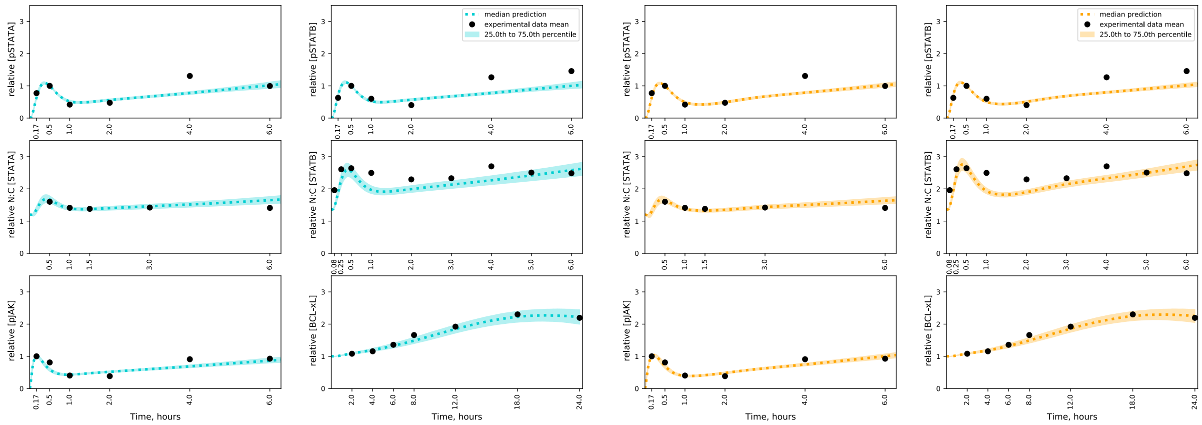


**Figure S9.** Predicted signaling responses derived by sampling either mode of *k24*. Dashed line, median prediction; shaded region; inter-quartile range; black dots, experimental predictions. Blue, predictions sampled from lower *k24* mode; orange, predictions sampled from upper *k24* mode.

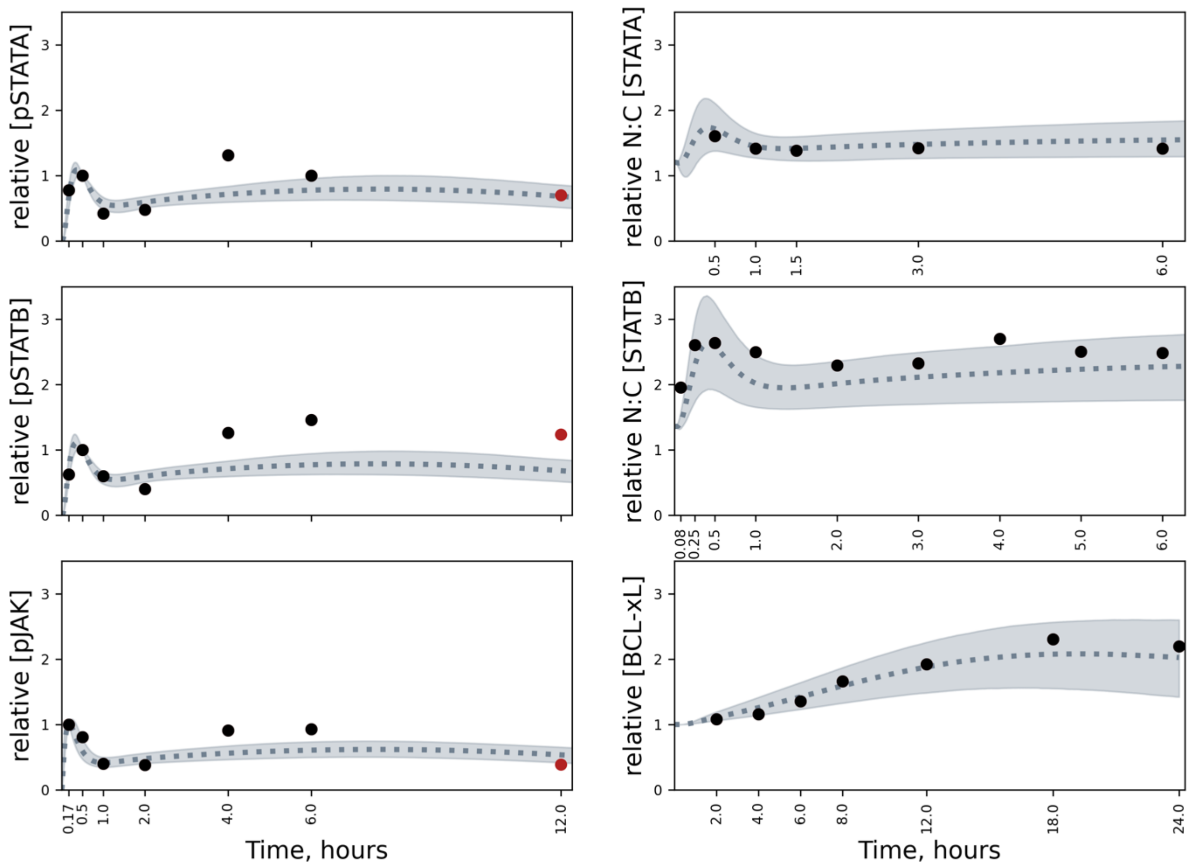


**Figure S10.** Model verification, extended data. Predicted signaling responses based on sampling the posterior distribution from the Bayesian parameter estimation approach. Dashed line, median model prediction; shading, the 5th to 95th percentiles of model predictions; black dots, baseline data experimental mean; red dots, extended time series experimental mean.

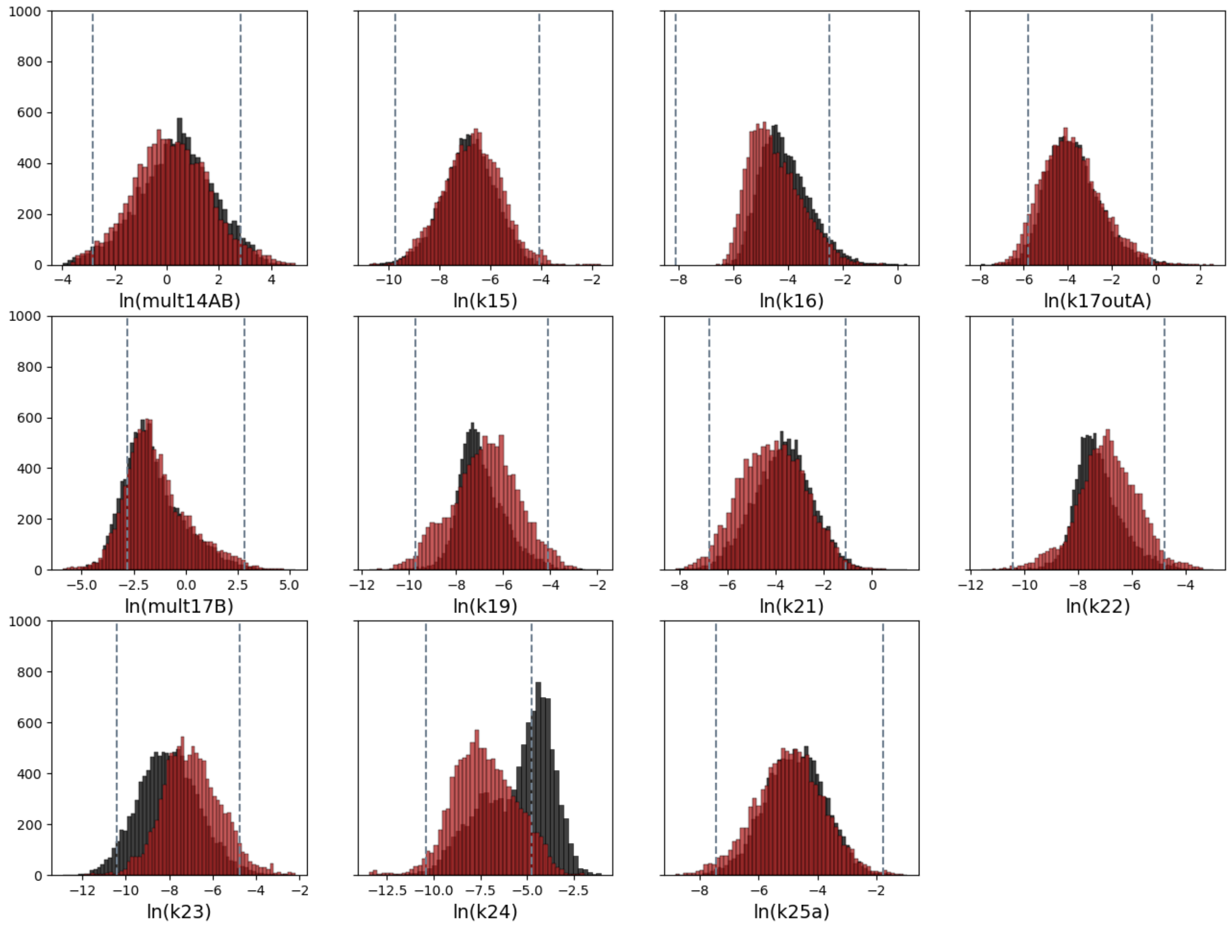


**Figure S11**. A representative sample of marginal posterior distributions before and after data integration, displayed on a natural log scale. Red, posterior estimated with extended data; black, posterior estimated with baseline data. Grey vertical lines, 2.5 to 97.5 percentiles of the prior distribution (which is not shown for clarity).
